# Coronavirus envelope protein drives iron sensing disorder by hijacking the TAp73-FDXR axis

**DOI:** 10.64898/2026.03.24.713916

**Authors:** Mingjun Zhu, Leyu Hu, Xingwei Fu, Boshui Yuan, Guiwen Guan, Lu Han, Zesheng Rong, Ru Tian, Gen Li, Muzi Du, Yixuan Ma, Ning Xu, Haixin Liu, Haolun Tian, Xinhuan Yin, Jianhui Zhong, Min Sun, Shanshan Yang, Shiyu Liu, Qingtao Liu, Jizong Li, Baochao Fan, Yiyang Chen, Qin Zhao, Ting Zhou, Lingling Chang, Xiaomin Zhao, Xin Ran, Qian Du, Siyuan Ding, Bin Li, Yong Huang, Dewen Tong

## Abstract

Iron overload is increasingly recognized as a critical contributor to coronavirus pathogenesis^1^, yet the underlying induction mechanisms remain unclear. Here, we uncover a fundamental pathway by which coronavirus drives IRP1 RNA-binding activity to induce iron accumulation^2^ via targeting the TAp73-FDXR axis. Specifically, coronavirus infection represses transcription of *FDXR* (encoding the key rate-limiting enzyme in host iron-sulfur cluster synthesis^3^), thereby impairing host iron-sulfur cluster generation to trigger the functional conversion of the cytosolic aconitase 1 (ACO1) into iron-regulatory protein 1 (IRP1)^4^, ultimately leading to the host’s persistently false perception of iron deficiency. We identify TAp73 as the primary transcription factor governing FDXR expression, and demonstrate that the coronavirus envelope protein (CoV-E) orchestrates TAp73 nuclear export. Subsequently, CoV-E binds TAp73 through a critical valine residue within its C-terminal PBM domain, inducing the K48-linked ubiquitination and proteasomal degradation of TAp73. Furthermore, we developed a CoV-E–targeting molecule, DPTP-FC, which blocks CoV-E–TAp73 interaction via forming steric hindrance and effectively alleviates iron accumulation and tissue damage caused by PEDV, PDCoV, and SARS-CoV-2 infection. Our study reveals the central role of the TAp73-FDXR axis in CoV-induced iron accumulation, highlighting CoV-E as an attractive antiviral target and DPTP-FC as a promising therapeutic candidate.

---

Iron is an essential trace element of nearly all living organisms^5^ that plays central roles in all kinds of life activities such as cell respiration^6^ and virus multiplication^7^. Given this absolute requirement, competition for iron between viruses and their hosts is inevitable. Nutritional immunity, a conserved innate defense mechanism in both vertebrates and invertebrates, describes the host’s strategy of restricting pathogen proliferation via limiting trace minerals (mainly iron) availability^8^. Nonetheless, certain pathogens subvert this defense by hijacking cellular iron through regulating iron metabolism, thereby antagonizing nutritional immunity to promote self-replication^9,10^. Coronaviruses (CoVs), as evolutionarily successful pathogens, have caused severe epidemics, represented by porcine epidemic diarrhea virus (PEDV, α), severe acute respiratory syndrome coronavirus 2 (SARS-CoV-2, β), infectious bronchitis virus (IBV, γ) and porcine deltacoronavirus (PDCoV, δ)^11^. Notably, previous research suggests that coronavirus infection is accompanied by iron overload^12,13^, while iron chelation can effectively ameliorate coronavirus-induced infection^14^. This suggests that iron dysregulation is a potential key factor in coronavirus pathogenesis, but the mechanisms by which coronavirus induces iron overload and the reason for the failure of the host’s iron homeostasis defense remain elusive.

Cellular iron homeostasis is maintained through the orchestrated regulation of iron metabolism, encompassing its transport, storage, and utilization^15^, all of which are masterfully governed by iron regulatory proteins (IRPs) IRP1 and IRP2^16^. IRPs modulate iron metabolism by binding to iron-responsive elements (IREs) in the 3’- or 5’-untranslated regions (UTRs) of target mRNAs, thereby stabilizing transcripts such as the transferrin receptor 1 (TfR1) or inhibiting the translation of others like ferroportin (FPN)^17^. However, it is unclear whether coronavirus-induced iron overload is mediated by manipulating IRPs. Notably, IRP1 is the iron-sulfur cluster (Fe-S, an ancient inorganic cofactor)-deficient form of cytosolic aconitase 1 (ACO1)^4^, and the stability of IRP2 is regulated by the E3 ubiquitin ligase adapter FBXL5, which itself requires Fe-S binding for its activity^18,19^. This convergence of evidence identifies Fe-S deficiency as a fundamental trigger for IRPs-mediated iron regulation. Fe-S biogenesis primarily occurs within the mitochondria^20,21^, and a critical initial step in this process is mediated by the ferredoxin reductase (FDXR), which transfers electrons from NADPH via the ferredoxin FDX2 to the core scaffold protein ISCU^22^. Previous studies have established that depletion of FDX2 or FDXR diminishes Fe-S clusters assembly and leads to mitochondrial iron overload^3,23^. Consequently, disorders in the synthesis of iron-sulfur clusters necessarily disrupted cellular iron homeostasis, but whether coronavirus precisely drives IRPs-mediated iron dysregulation by targeting host Fe-S biogenesis remains entirely unexplored.

## Iron accumulation boosts RdRp activity

We first confirmed that CoVs, including PEDV and SARS-CoV-2, consistently induce cellular iron accumulation (Extended Data Fig. 1a,b). Elevated iron levels significantly promoted viral replication, whereas iron chelation suppressed it (Extended Data Fig. 1c-f). Accordingly, iron supplementation exacerbates SARS-CoV-2 and PEDV infection, increasing viral load (Extended Data Fig. 1g,h) and tissue damage (Extended Data Fig. 1i,j), and reducing survival or weight gain in both piglets and humanized ACE2 mice (Extended Data Fig. 1k,l). These findings establish iron accumulation as a key factor in CoVs pathogenesis. Previous studies suggested that SARS-CoV-2 RdRp harbors Fe-S, metal cofactors critical for viral replication^24^, and that zinc can replace endogenous Fe-S under aerobic conditions^25,26^. Whereafter, we discovered that cellular iron significantly enhanced the RNA-dependent RNA polymerase (RdRp, also called nsp12) activity of PEDV (Extended Data Fig. 2a). Through subsequent analysis and anaerobic purification after eukaryotic expression, we identified a conserved zinc-finger binding motif in CoVs-RdRp, comprising two catalytic domains H272S-C278-C283-C287 and C464-H619-C622-C623 (Extended Data Fig. 2b,c). Consistent with previous evidence, mutating either domain (by replacing any set of three Cys residues with Ser) exhibited a reduction of half or even more shoulder at ∼420 nm in its ultraviolet-visible (UV-vis) absorption spectrum (Extended Data Fig. 2d,e), supporting the presence of Fe-S coordinated by these motifs. Given that the LYR like motifs is recognized by the cytosolic iron-sulfur cluster assembly (CIA) machinery and also interact with the Fe-S biogenesis cochaperone HSC20 (also known as HSCB) and so on^27^. We also analyzed RdRp sequences across CoVs and identified conserved and strain-specific LYR-like motifs (Extended Data Fig. 2f). Homology modeling revealed PEDV RdRp spatial placement, and subsequent mutagenesis of two motifs (VYR and LYK, but not AFK) to triple alanine (AAA) decreased 420 nm absorbance (Extended Data Fig. 2g-i), confirming their role in Fe-S coordination. Co-immunoprecipitation (Co-IP) and confocal laser scanning microscopy (CLSM) confirmed that RdRp interacts with HSC20, as well as the chaperone HSPA9 and CIAO1, while substitution of either of the two LYR motifs (VYR and LYK) with AAA reduced HSC20 binding (Extended Data Fig. 2j,k). The above evidence collectively indicates that Fe-S and their assembly mechanism are conserved in CoVs-RdRp. Using TEMPOL (a specific dissociative agent of Fe-S^24^), we demonstrated that iron enhances CoVs-RdRp activity in a Fe-S–dependent manner (Extended Data Fig. 2l-n).

## CoVs infection activates IRPs

To elucidate the mechanism by which CoVs induce intracellular iron accumulation, we first investigated the impact of CoVs infection on cellular iron metabolism. Our data indicate that PEDV infection exerts broad effects on cellular iron metabolism, resulting in a marked upregulation of TfR1 and simultaneous downregulation of FPN (Fig. 1a), which can collectively mediate a global increase in cellular iron levels. As previously mentioned, IRPs serve as central regulators of cellular iron homeostasis, with their activation tightly dependent on the assembly status of host Fe-S clusters^16,28^. Notably, IRP1 can be directly activated upon loss of the Fe-S clusters from ACO1 (Fig. 1b)^4,29^. Coincidentally, PEDV infection didn’t affect IRP1 expression but markedly reduced ACO1 enzymatic activity (Fig. 1c,d). We therefore hypothesized that PEDV infection induces Fe-S loss in ACO1, thereby converting it into IRP1. To test this, we designed a fluorescent probe-labeled nucleic acid aptamer (Cy5-lebeled NAA) that specifically binds IRP1 and assessed its RNA-binding activity using electrophoretic mobility shift assay (EMSA) (Fig.1e,f). As expected, ACO1/IRP1, enriched anoxically following PEDV infection (Fig.1g), exhibits marked increase in RNA-binding activity (Fig. 1h,i). Concurrently, we detected a pronounced reduction in absorbance at ∼420 nm for ACO1/IRP1 (Fig. 1j), indicating that PEDV infection triggers a functional conversion from ACO1 to IRP1 via loss of its Fe-S cluster.

**Fig. 1:**
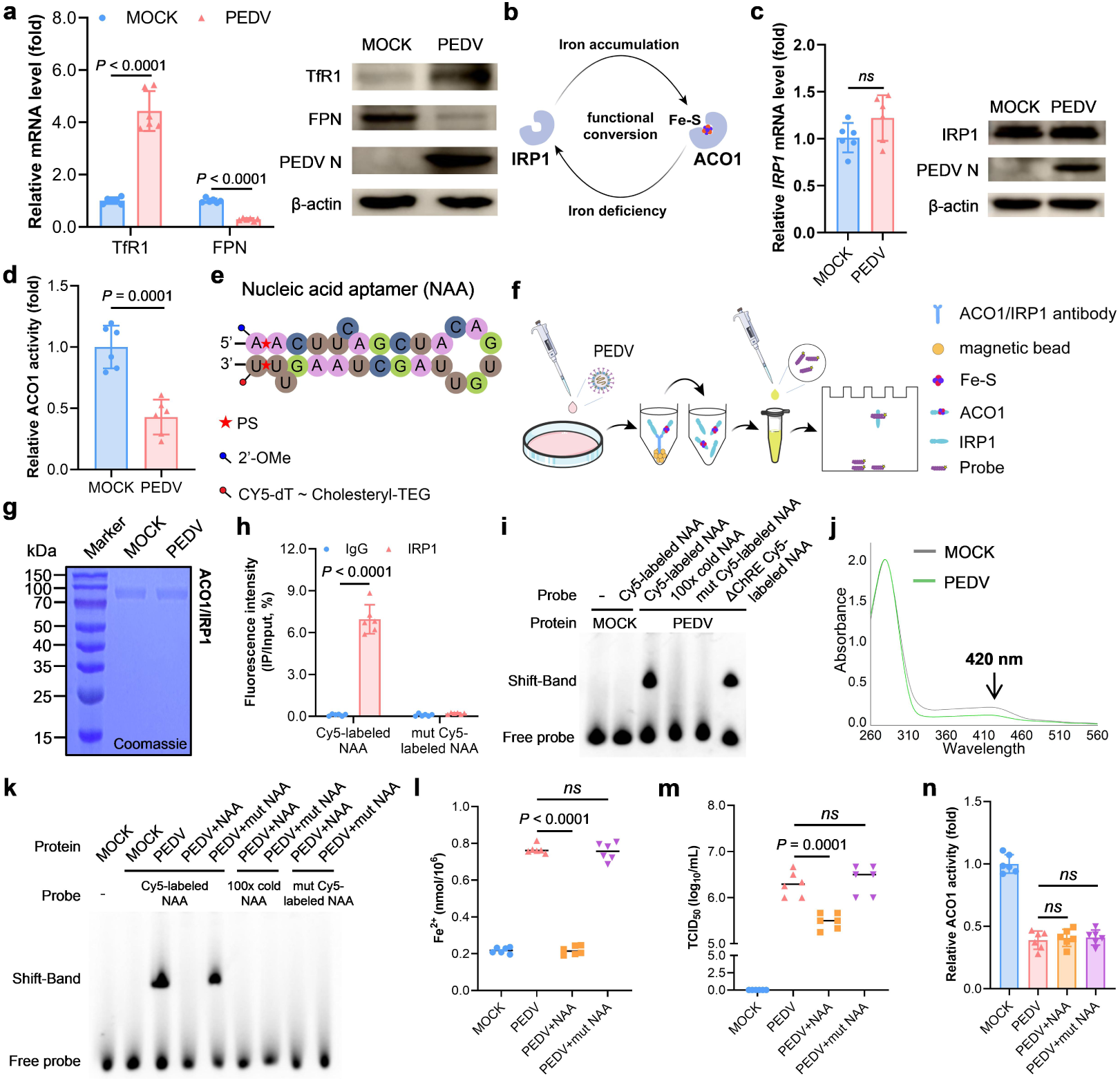
PEDV infection disrupts IRP1 function. **a-b**, Relative mRNA levels and western blot analysis of Trf1 and FPN (**a**) and IRP1 (**b**) in IPEC-J2 cells with or without PEDV infection for 16 h (*n*=6). **c**, Schematic diagram of the functional conversion between IRP1 and ACO1. **d**, ACO1 activity in IPEC-J2 cells with or without PEDV infection for 16 h (*n*=6). **e**, Schematic diagram of nucleic acid aptamer (NAA) sequence and modification. **f**, Experimental setup for evaluation of the RNA binding activity of IRP1. **g**, Coomassie Brilliant Blue staining of anaerobically purified ACO1/IRP1 from mock and PEDV-infected IPEC-J2 cells. **h**, RIP analysis of the specific binding of IRP1 proteins to NAA probes. **i**, EMSA of the specific binding of IRP1 proteins to NAA probes. **j**, UV-Vis absorption spectra of anaerobically purified ACO1/IRP1 from mock and PEDV-infected IPEC-J2 cells. **k**, EMSA of IRP1 binding to NAA and mut NAA probes in IPEC-J2 cells. **l-n**, Fe²⁺ levels (**l**), virus titer (**m**), and ACO1 activity (**n**) of IPEC-J2 cells treated with NAA or mut NAA for 16 h (*n*=6).Data are represented as mean±SD and analyzed by two-tailed Student’s t-test (**a, b, d, h**) or one-way ANOVA with Tukey’s multiple comparisons test (**l-n**).

Meanwhile, NAA can also act as a blocker of the RNA binding activity of IRP1. We found that NAA treatment effectively reversed the enhanced IRP1 RNA-binding activity induced by PEDV infection (Fig. 1k), subsequently restoring the PEDV-induced cellular iron accumulation and potently suppressed viral replication (Fig. 1l,m), indicating that the RNA binding activity of IRP1 is a key factor for CoVs-induced iron accumulation promoting its own replication. Meanwhile, NAA did not affect ACO1 activity (Fig. 1n), confirming that the observed effects are specifically mediated by IRP1. Collectively, these results define a mechanism whereby CoVs infection activates IRP1 by obstructing host Fe-S synthesis.

## FDXR suppression underlies IRP1 activation

Integrated transcriptomic analysis of PEDV-infected IPEC-J2 cells revealed that differentially expressed genes (DEGs) were predominantly enriched in innate immune pathways, indicating a robust suppressive effect on host antiviral immunity (Extended Data Fig. 3a). This observation aligns with established evidence that iron overload can precipitate systemic immunosuppression^10^. To further delineate the metabolic impact of infection, we prioritized 103 iron metabolism-related DEGs for protein-protein interaction (PPI) network analysis, which identified 8 distinct functional modules (Fig. 2a and Extended Data Fig. 3b). A prominent core network, comprising 13 genes dedicated to iron-sulfur (Fe-S) cluster biogenesis, was found to be centered on FDXR (Fig. 2b). As an essential electron transporter and a core regulatory component of the Fe-S cluster synthesis pathway, the transcriptional level of FDXR was significantly downregulated following PEDV infection, identifying it as a pivotal node in the virus-induced metabolic dysregulation (Fig. 2c,d). We then confirmed that FDXR expression was inhibited by PEDV infection in IPEC-J2 cells (Extended Data Fig. 4a,b). Consistent with this, FDXR was significantly down-regulated in all three segments of the small intestine exhibiting the most robust PEDV replication (Extended Data Fig. 4c-e). Further evidence demonstrated that FDXR overexpression mitigates iron accumulation induced by PEDV, whereas FDXR knockout leads to a further escalation in iron accumulation regardless of infection status (Extended Data Fig. 4f,g), suggesting that FDXR serves as a crucial regulator of PEDV-induced iron accumulation. Correspondingly, PEDV replication was inhibited by FDXR overexpression and promoted by FDXR knockout (Extended Data Fig. 5a-h). Importantly, it seems to be a conserved feature among CoVs, as FDXR knockdown similarly enhanced cellular iron levels and SARS-CoV-2 replication in HeLa-ACE2 cells (Extended Data Fig. 6a-d). The poly-U template catalysis by the nsp12-nsp7-nsp8 complex, harvested under hFDXR-knockdown conditions, showed intensive RNA polymerization activity, which was effectively inhibited by complementary treatment with the iron chelator Dp44mT (Extended Data Fig. 6e-g). Meanwhile, the iron accumulation caused by FDXR knockdown was alleviated by Dp44mT treatment (Extended Data Fig. 6h). Furthermore, RdRp purified anoxically from hFDXR-knockdown Expi293F cells exhibited enhanced absorbance at 420 nm relative to wild-type cells, which was reduced upon Dp44mT treatment (Extended Data Fig. 6i,j), suggesting the crucial role of FDXR in mediating the coronavirus-induced iron accumulation that enhances RdRp activity. Despite RdRp’s inherent lack of proofreading capability, we found that cellular iron which potentiates RdRp activity did not elevate the frequency of defective viral particles (Extended Data Fig. 6k). It may be attributed to the combined effects of nsp7/nsp8-mediated template strand stabilization and the error-correcting exonuclease activity of nsp14^30–32^.

**Fig. 2:**
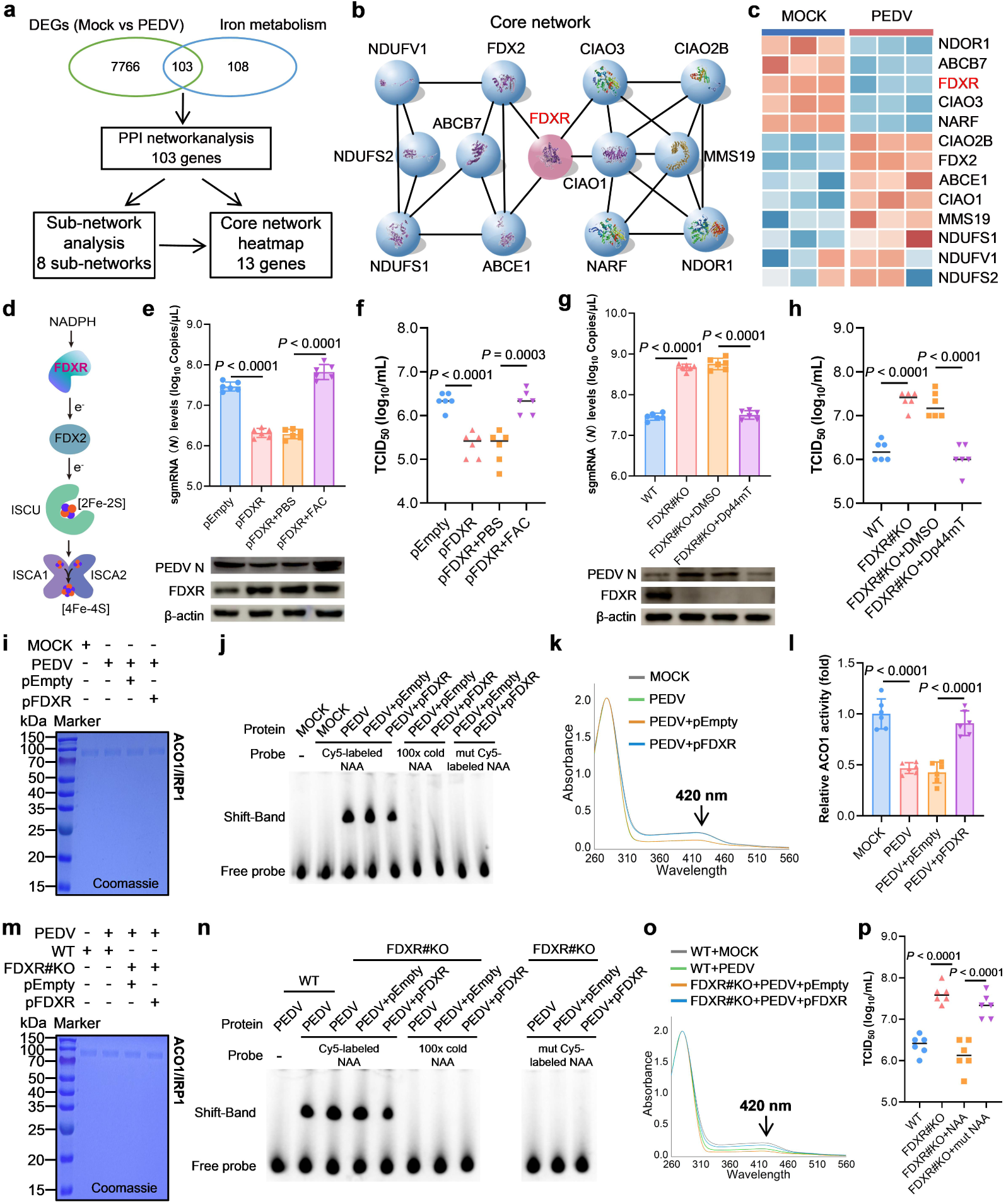
FDXR suppresses PEDV replication through IRP1 regulation by Fe-S assembly. **a**, Schematic workflow for identifying iron metabolism-related differentially expressed genes (DEGs) between Mock and PEDV-infected cells. The overlap between DEGs and iron metabolism genes was used for Protein-protein interaction (PPI) network construction, followed by sub-network analysis and core network heatmap generation. **b**, PPI network of Fe-S cluster biosynthesis and assembly-related proteins. FDXR is highlighted in red, indicating its central role in the core network. **c**, Heatmap showing transcriptional changes of core Fe-S cluster biosynthesis and assembly genes upon PEDV infection (Mock vs PEDV). **d**, Schematic diagram of Fe-S assembly mediated by FDXR. **e, f**, Absolute quantitative, western blot (**e**) and virus titer (**f**) analysis of pEmpty or pFDXR-transfected IPEC-J2 cells in the presence or absence of FAC for 16 h (*n* = 6). **g, h**, Absolute quantitative, western blot (**g**) and virus titer (**h**) of IPEC-J2 cells and FDXR#KO IPEC-J2 cells in the presence or absence of Dp44mT for 16 h (*n* = 6). **i-k**, Coomassie Brilliant Blue staining (**i**), EMSA (**j**) and UV-Vis absorption spectra (**k**) of anaerobically purified ACO1/IRP1 in mock and PEDV infected Expi293F cells with pEmpty or pFDXR-transfected. **l**, ACO1 activity of IPEC-J2 cells, and PEDV infected IPEC-J2 cells with Empty or FDXR transfection for 16 h (*n* = 6). **m-o**, Coomassie Brilliant Blue staining (**m**), EMSA (**n**) and UV-Vis absorption spectra (**o**) of anaerobically purified ACO1/IRP1 in mock and PEDV infected Expi293F cells, PEDV infected FDXR#KO Expi293F cells with pEmpty or pFDXR-transfected. **p**, Virus titer of PEDV infected IPEC-J2 cells and FDXR#KO IPEC-J2 cells with NAA or mut NAA treatment for 16 h (*n* = 6). Data are represented as mean ± SD, and analyzed by two-tailed Student’s t-test (**f-h, l, p**).

To determine whether the effect of FDXR on CoVs replication depends on its regulation of cellular iron metabolism, we conducted a reverse validation of the impact of FDXR on CoVs replication and demonstrated that PEDV replication was significantly suppressed with FDXR overexpression and reversed after complementing with FAC treatment, but promoted by FDXR-knockout and reversed through Dp44mT treatment (Fig. 2e-h), indicating that FDXR involved CoVs replication in a manner dependent on its regulation of cellular iron metabolism. Building on these observations, we next investigated whether FDXR mediates the PEDV-driven functional conversion from ACO1 to IRP1, a transition that triggers a cellular misperception of iron deficiency. Our results showed that FDXR overexpression significantly attenuated PEDV-induced RNA-binding activity of IRP1 (Fig. 2i,j); correspondingly, Fe-S cluster assembly on ACO1 was markedly increased, accompanied by enhanced aconitase activity (Fig. 2k,l). Consistently, FDXR knockout further intensified PEDV-induced IRP1 activation and exacerbated the loss of Fe-S in ACO1, but both phenotypes can be reversed by supplementing FDXR expression. (Fig. 2m-o). Furthermore, blocking IRP1 RNA-binding activity with NAA abolished the effect of FDXR knockout on PEDV replication (Fig. 2p). Collectively, these findings demonstrate that PEDV facilitates RdRp activity to promote self-replication by downregulating FDXR to drive sustained IRP1 activation, thereby orchestrating the cellular iron accumulation

## TAp73 mediates CoV suppression of FDXR

To elucidate the mechanism by which CoV downregulates FDXR, we employed actinomycin D (ActD) treatment following PEDV infection (Extended Data Fig. 7a). Our findings confirmed that PEDV infection exerts no significant effect on *FDXR* mRNA stability and post-transcriptional process, indicating that the suppression occurs at *FDXR* transcription (Extended Data Fig. 7b-d). To identify the transcriptional regulators implicated in PEDV-mediated repression of *FDXR* transcription, we constructed a luciferase reporter under the control of 2,000 base pairs (bp) of the porcine *FDXR* promoter sequences, and determined the core promoter region of FDXR ranges from −700 to −1 (M3) (Extended Data Fig. 8a). Bioinformatic screening of the core promoter using the JASPAR vertebrate database (http://jaspar.genereg.net/)^33^ identified multiple putative transcription factor binding sites (TFBS) including p53 family members (TAp53, TAp63, and TAp73), as well as for TBX1, ZEB1, PRDM1, GBX2, and SNAI1 (Extended Data Fig. 8b). Dual-luciferase reporter gene assay confirmed that TAp73, rather than other transcription factors, mediates the activation of *FDXR* promoter (Extended Data Fig. 8c). Consistently, chromatin immunoprecipitation (ChIP) assay showed that TAp73 directly bound to the *FDXR* promoter region (Extended Data Fig. 8d,e). Furthermore, both the overexpression and knockout of TAp73 confirmed its role as a critical transcriptional activator of FDXR and established its necessity in regulating cellular iron homeostasis (Extended Data Fig. 8f-i).

To determine whether TAp73 serves as the primary mediator of PEDV-induced FDXR transcriptional suppression, we initially assessed its expression profiles following viral infection. We observed that PEDV significantly downregulated TAp73 protein levels without affecting its mRNA abundance, suggesting that TAp73 is regulated post-transcriptionally during infection (Fig. 3a). To further elucidate this mechanism, we utilized the methionine analogues L-homopropargylglycine (HPG) in conjunction with click chemistry^34^ to monitor nascent TAp73 protein synthesis (Fig. 3b). The results showed that PEDV infection had no discernible effect on TAp73 synthesis (Fig. 3c), indicating that the reduction in TAp73 levels is driven by enhanced protein degradation. Notably, the dose-dependent restoration of TAp73 effectively rescued FDXR transcription (Fig. 3d) and significantly mitigated both intracellular iron accumulation (Fig. 3e) and the elevated IRP1 RNA-binding activity (Fig. 3f). Collectively, these findings establish TAp73 as a critical regulatory hub through which PEDV suppresses FDXR expression to trigger iron accumulation. To further dissect the hierarchy of the TAp73-FDXR axis and its role in CoVs-induced iron accumulation, we characterized the phenotypic changes of TAp73 or FDXR reconstitution in their respective knockout (KO) backgrounds. TAp73-KO not only exacerbated the iron accumulation caused by PEDV (Fig. 3g), amplified IRP1 RNA-binding activity (Fig. 3h) and significantly enhanced viral replication (Fig. 3i,j). Replenishing either TAp73 or FDXR effectively mitigated these pathological changes, among which FDXR restoration yielded a more pronounced rescue effect (Fig. 3g-j). Consistent with its role as a potential downstream effector, FDXR-KO phenocopied the effects of TAp73 deficiency, leading to further iron accumulation (Fig. 3k), hyperactivated IRP1 (Fig. 3l) and elevated viral titers (Fig. 3m,n). Crucially, while FDXR add-back entirely reversed these phenotypes in FDXR-KO cells, the restoration of TAp73 failed to yield any significant improvement (Fig. 3k-n). These data unequivocally establish FDXR as a downstream mediator of TAp73, identifying the TAp73-FDXR axis as a pivotal pathway through which CoVs activate IRP1 to drive intracellular iron accumulation.

**Fig. 3:**
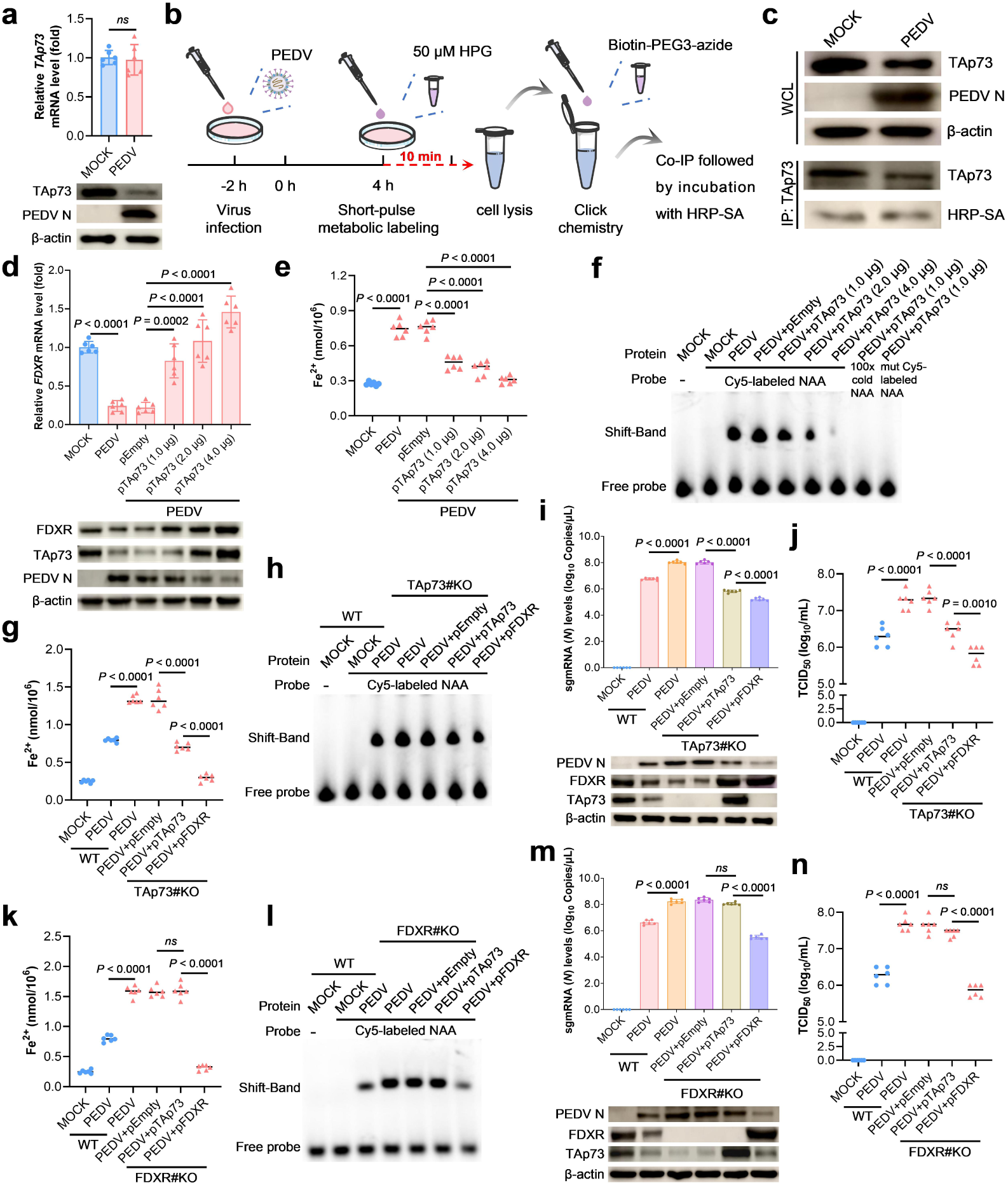
TAp73 regulates cellular iron homeostasis through the FDXR-IRP1 ax is to inhibit PEDV replication. **a**, Relative mRNA levels and western blot an alysis of IPEC-J2 cells with or without PEDV infection for 16 h (*n*=6). **b-c**, E xperimental setup (**b**) and IP analysis (**c**) for evaluating the binding activity of TAp73 with new protein. **d-f**, Relative levels, western blot (**d**), Fe²⁺ levels (**e**), and EMSA (**f**) of PEDV-infected IPEC-J2 cells with or without TAp73 dose-d ependent transfection for 16 h (*n*=6). **g-j**, Fe²⁺ levels (**g**), EMSA (**h**), relative l evels and western blot (**i**), and virus titer (**j**) of PEDV-infected IPEC-J2 cells a nd TAp73#KO IPEC-J2 cells with or without TAp73/FDXR transfection for 16 h (*n*=6). **k-n**, Fe²⁺ levels (**k**), EMSA (**l**), relative levels and western blot (**m**), and virus titer (**n**) of PEDV-infected IPEC-J2 cells and FDXR#KO IPEC-J2 c ells with or without TAp73/FDXR transfection for 16 h (*n*=6).Data are represen ted as mean±SD and analyzed by two-tailed Student’s t-test (**a**) or one-way A NOVA with Tukey’s multiple comparisons test (**d**, **e**, **g**, **i-k**, **m**, **n**).

## CoV-E protein degrades TAp73

To further elucidate the mechanism by which PEDV degraded TAp73, we generated wild-type viruses and UV-inactivated viruses to examine whether this process is associated with viral structural proteins or non-structural proteins. We found that PEDV, even after UV-inactivation, still significantly repressed the TAP73-FDXR axis, indicating the crucial role of the structural proteins (Fig. 4a). We subsequently constructed eukaryotic expression vectors of spike protein (S), envelope protein (E), membrane protein (M) and nucleocapsid protein (N) of PEDV and demonstrated that PEDV E protein is the key factor that suppresses TAp73-FDXR axis (Fig. 4b). Existing evidence indicated that E proteins of CoVs interact with host proteins through its C-terminal PDZ-binding motif (PBM) and exerts various regulatory functions^35,36^. We examined that the C-terminal region of the E protein for different subtypes PEDV contains a conserved PBM domain and the hydrophobic valine (V) residues at the tail end which provide a structural basis for its interaction with host proteins through hydrophobic forces (Fig. 4c). To verify the function of this residue, we constructed the mutant plasmid pPEDV E (V76G) and the recombinant strain rPEDV-E/V76G by substituting hydrophobic Val76 with hydrophilic glycine (G) and demonstrated that the valine in PDM domain of E is crucial for PEDV E regulating the TAp73-FDXR axis (Fig. 4d,e). Furthermore, we also found that the valine in PBM domain of the E protein among different genera of the CoVs remained relatively conserved, with only IBV (γ genus) having differences (Extended Data Fig. 9a). Consistent with this observation, both SARS-CoV-2 E and PDCoV E can also regulate TAp73-FDXR axis but out of action by substituting V with G (Extended Data Fig. 9b,c). However, IBV E functioned weakly on the TAp73-FDXR axis, but enhanced miraculously by substituting threonine (T) with V, which might be because the T residue in PBM domain weaken its hydrophobicity (Extended Data Fig. 9d). The above research findings suggested that the V residue at the C-terminus of the E protein plays a crucial role in modulating the TAp73-FDXR axis.

**Fig. 4:**
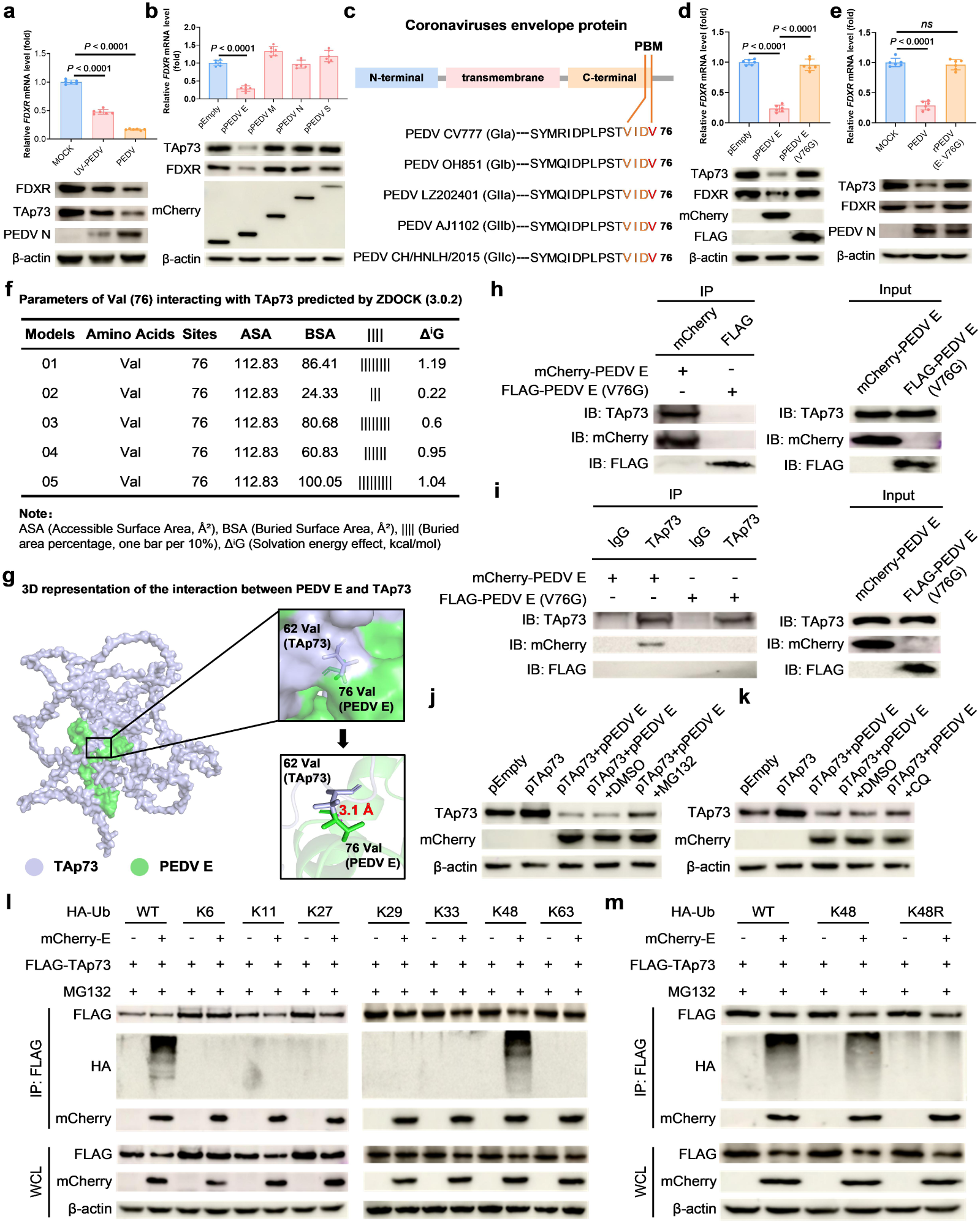
CoVs E suppresses *FDXR* transcription by mediating the K48-linked ubiquitination degradation of TAp73. **a**, Relative mRNA levels and western blot analysis of analysis of IPEC-J2 cells with PEDV and Ultraviolet-inactivated PEDV (UV-PEDV) infection for 16 h (*n* = 6). **b**, Relative mRNA levels and western blot analysis of analysis of IPEC-J2 cells transfected PEDV structural proteins for 16 h (*n* = 6). **c**, Domain analysis of PEDV E proteins of each subtype. **d**, Relative mRNA levels and western blot analysis of analysis of IPEC-J2 cells with PEDV E or PEDV E (V76G) transfection for 16 h (*n* = 6). **e**, Relative mRNA levels and western blot analysis of analysis of IPEC-J2 cells with PEDV or rPEDV (E: V76G) infection for 16 h (*n* = 6). **f**, Predicted interaction parameters of Val76 with TAp73 generated using ZDOCK (v3.0.2). **g**, Three-dimensional representation of the interaction between PEDV E protein and TAp73. TAp73 is shown in gray and PEDV E protein in green; key interacting residues are indicated. **h, i**, Co-IP analysis of the interaction between TAp73 and PEDV E or PEDV E (V76G) in IPEC-J2 cells. **j, k**, Western blot analysis of IPEC-J2 cells transfected TAp73 with or without PEDV E for 16 h, followed by MG132 (**j**) or CQ (**k**) treatment for 6 h. **l, m**, IP analysis of the ubiquitination modification type of PEDV E protein degrading TAp73 in HEK293T cells. Data are represented as mean ± SD, and analyzed by one-way ANOVA with Tukey’s multiple comparisons test (**a, b, d, e**).

Subsequently, we investigated whether CoV-E interact with TAp73 and whether V is the key factor to this interaction between them. The docked protein models of TAp73 and PEDV E were constructed for preliminary evaluation and screening, with a focus on Val76 of the PEDV E. Five prediction models were listed, among which model 05 exhibited the highest buried surface area (BSA) value (Fig. 4f). As shown in Fig. 4g, we conducted homology modeling based on the crystal structure of CoV E (PDB ID: 2MM4) and the predicted structure of TAp73 (Alphaphold DB ID: O15350) to produce a more exact model of this region. Three-dimensional spatial distance calculations revealed that the Val76 of the E protein and Val62 of TAp73 are in close proximity, with a distance of 3.1 Å, which aligns with the typical criteria for hydrophobic interactions (generally < 5 Å)^37^. In contrast to the conserved Val62 in TAp73, TAp53 and TAp63 possess a highly hydrophilic aspartic acid at the equivalent position, potentially precluding similar hydrophobic interactions with the E protein (Extended Data Fig. 9e). Next, we verified the interaction between these two proteins using co-immunoprecipitation (Co-IP) assays and confirmed that valine residue within the PBM domain is essential for this interaction (Fig. 4h,i), a finding consistent with earlier theoretical predictions. It is generally believed that CoV-E are located in the endoplasmic reticulum, while TAp73 is located in the nucleus^38,39^. Strangely, how does the interaction between them occur? We unexpectedly discovered that PEDV E mediates the nuclear export of TAp73 (Extended Data Fig. 10a-d) and found this process required for PEDV- induced degradation of TAp73 (Extended Data Fig. 10e,f). The ubiquitin-proteasome system and the autophagy-lysosomal system are two major clearance systems responsible for intracellular homeostasis^40^. Ulteriorly, we found that CoV-E mediate TAp73 degradation via the ubiquitin-proteasome system rather than autophagy-lysosomal system (Fig. 4j,k). Considering that different types of polyubiquitination modifications, such as those linked to K6, K11, K27, K29, K33, K48 and K63, are involved in regulating the functions of cellular proteins^41^, we systematically investigate which type of ubiquitination modification is closely associated with the degradation of TAp73 mediated by CoV-E. The results showed that CoV-E specifically mediate the K48-linked polyubiquitination of TAp73 (Fig. 4l,m). Taken together, CoV-E suppress *FDXR* transcription by mediating the nuclear export and K48-linked ubiquitination and proteasomal degradation of TAp73.

## DPTP-FC blocks CoV by targeting E

Virtual screening which is a vital computational strategy for identifying lead compounds from large databases performed to identify potential small-molecule inhibitors targeting Val76 of the PEDV E protein. A comprehensive screening of 99,288 compounds from the Diverse-lib database was conducted using the Lipinski filter. Thus, 1,500 candidates have been identified with binding scores ranging from -6.3 to -8.3 kcal/mol (Supplementary Data Set). Due to the high binding scores of -7.8 kcal/mol to -8.3 kcal/mol, respectively, five compounds identified as CID661823, CID16673644, CID5307372, CID16191144, CID4147703 were selected for further analysis (Extended Table 1). Among the top hits, an amide-type compound (named DPTP-FC) exhibited potential to form superior spatial steric hindrance and was therefore selected for further experimental validation (Extended Data Fig. 11a,b). Next, the DPTP-FC were synthesized via the amidation reaction between amino and carboxylic acid groups (Fig. 5a). Its chemical structure was comprehensively characterized and confirmed by ^1^H NMR, ^13^C NMR, HRMS and UV-vis spectra. (Extended Data Fig. 12a-d). We subsequently demonstrated that DPTP-FC exerted low micromolar antiviral activity against a broad panel of CoVs (including PEDV, SARS-CoV-2, and PDCoV, but was inactive against IBV) with half-maximal effective concentration (EC_50_) of 581.2 nM, 793.7 nM and 1.087 μM, respectively (Fig. 5b,c and Extended Data Fig. 13a,b). Notably, the anti-PEDV potency of DPTP-FC surpassed that of Remdesivir (EC_50_ values of 1.651 μM), although its efficacy against SARS-CoV-2 was inferior to Remdesivir (EC_50_ values of 98.07 nM) (Fig. 5d,e). We then demonstrated that DPTP-FC effectively disrupted the PPI between PEDV E and TAp73 (Fig. 5f). This intervention successfully rescued the FDXR expression suppressed by PEDV infection and restored the diminished ACO1 enzymatic activity (Fig. 5g,h). Crucially, DPTP-FC treatment abrogated the PEDV-induced elevation of IRP1 RNA-binding activity while promoting the re-assembly of the Fe-S cluster in the ACO1/IRP1 complex (Fig. 5i-k). These findings identify DPTP-FC as a potent small-molecule inhibitor capable of reversing the PEDV-induced functional conversion from ACO1 to IRP1, thereby maintaining cellular iron homeostasis and metabolic stability during viral challenge.

**Fig. 5:**
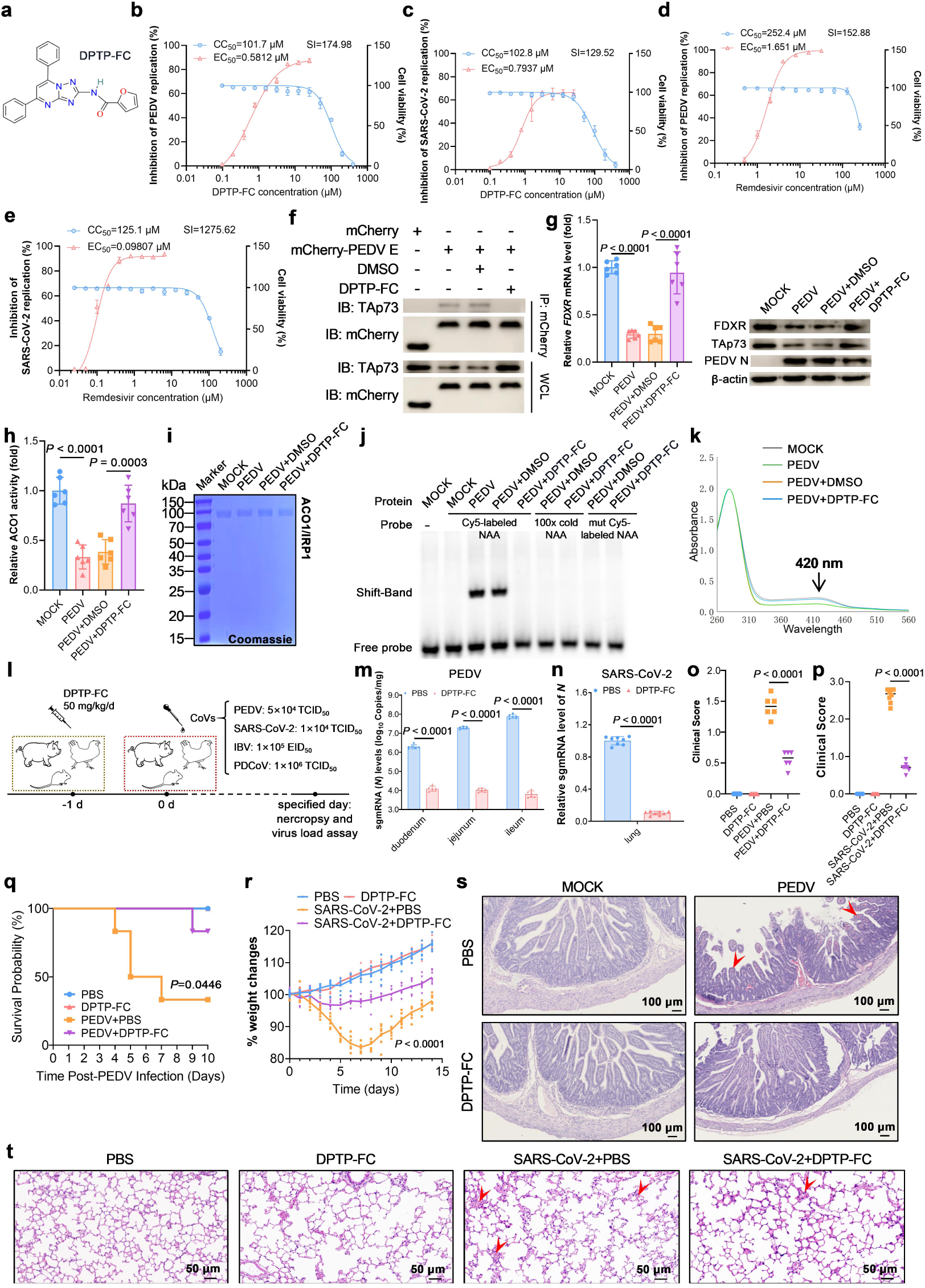
DPTP-FC inhibits CoVs replication by targeting the TAp73-E interaction to suppressing the TAp73-FDXR-IRP1 axis. **a**, Chemical structure of DPTP-FC. **b, c**, The half-maximal effective concentration (EC_50_) of DPTP-FC (**b**) and remdesivir (**c**) against PEDV, together with the 50% cytotoxic concentration (CC_50_) values, as measured in IPEC-J2 cells. SI=CC_50_/EC_50_. **d, e**, The EC_50_ of DPTP-FC (**d**) and remdesivir (**e**) against SARS-CoV-2, together with the CC_50_ values, as measured in HeLa-ACE2 cells. **f-h**, IP analysis (**f**), relative levels and western blot analysis (**g**), and ACO1 activity (**h**) in PEDV-infected IPEC-J2 cells in the presence or absence of DPTP-FC. **i-k**, Coomassie Brilliant Blue staining (**i**), EMSA (**j**), and UV-Vis absorption spectra (**k**) of anaerobically purified ACO1/IRP1 from PEDV-infected Expi293F cells in the presence or absence of DPTP-FC. **l**, Experimental setup for evaluating the in vivo anti-coronavirus activity of DPTP-FC. **m-t**, Effects of DPTP-FC treatment on PEDV and SARS-CoV-2 infection in vivo.In PEDV LZ202401-infected piglets, PEDV N gene copy numbers in intestinal tissues (**m**), clinical scores of small-molecule-treated and control piglets, as well as infected piglets with or without DPTP-FC treatment (**o**), survival curves (**q**), and H&E staining of jejunal tissues (**s**), with representative pathological lesions indicated by red arrows (*n*=6). In SARS-CoV-2 SZTH-003-infected hACE2-transgenic mice, relative viral RNA levels in lung tissues (**n**), clinical scores of small-molecule-treated and control mice, as well as infected mice with or without DPTP-FC treatment (**p**), percentage body weight changes (**r**), and H&E staining of lung tissues (**t**), with representative pathological lesions indicated by red arrows (*n*=8). Data are presented as mean±SD and were analysed by two-tailed Student’s t-test (**g, h, m-o, p, r**) or the Gehan-Breslow-Wilcoxon test (**q**).

Subsequently, the efficacy of DPTP-FC was assessed in piglets, chicks and humanized ACE2 mice via daily injections at a dosage of 50 mg per kg (Fig. 5l). DPTP-FC treatment resulted in a 2∼3log_10_ reduction in PEDV viral RNA titers and an ∼85% decrease in SARS-CoV-2 RNA levels (Fig. 5m,n). Clinical scoring revealed that DPTP-FC administration significantly ameliorated clinical symptoms induced by PEDV, SARS-CoV-2, and PDCoV, whereas it had no discernible effect on IBV-infected chicks (Fig. 5o,p and Extended Data Fig. 13c,d). Specifically, DPTP-FC significantly enhanced the survival rates of PEDV- and PDCoV-infected piglets and normalized weight recovery in SARS-CoV-2-infected mice (Fig. 5q,r and Extended Data Fig. 13e), but failed to improve weight gain in IBV-infected chicks (Extended Data Fig. 13f). Histopathological assessment further confirmed that DPTP-FC attenuated tissue damage in the intestines or lungs following PEDV, SARS-CoV-2, and PDCoV challenge (Fig. 5s,t and Extended Data Fig. 13g), but was ineffective against IBV-induced tracheal epithelial lesions (Extended Data Fig. 13h). Taken together, these findings identify DPTP-FC as a promising small-molecule drug candidate that exerts its antiviral efficacy by effectively thwarting the viral hijacking of cellular iron metabolism. Importantly, our work provides a novel therapeutic perspective, demonstrating that restoring host iron homeostasis represents a viable and potent strategy for antiviral intervention. This study underscores the potential of targeting virus-induced metabolic dysregulation as a paradigm for developing next-generation broad-spectrum antivirals.

## Discussion

While the host innate immune system employs nutritional immunity as a crucial mechanism to restrict iron availability for invading microbes^8,42^, successful pathogens antagonize nutritional immunity by utilizing a wide range of strategies to plunder cellular iron^43^. CoVs infection are frequently characterized by cellular iron overload^1^, however, the mechanisms preventing the host’s homeostatic machinery from correcting this dysregulation have remained elusive. Here, we uncover a global manipulative strategy by which CoVs subvert cellular iron metabolism. CoVs exploit the E protein to induce TAp73 degradation, thereby suppressing FDXR transcription and abrogating host Fe-S cluster biogenesis^3^. This deficiency triggers the functional transition of ACO1 into IRP1^44^, as well as the activation of IRP2^18,19^, effectively locking the sensor in a state of perceived iron deficiency. The resulting persistent activation of IRP1 drives a cascade of pathological iron accumulation (Fig. 6), providing a mechanistic explanation for the functional failure of host iron homeostasis during infection.

**Fig. 6:**
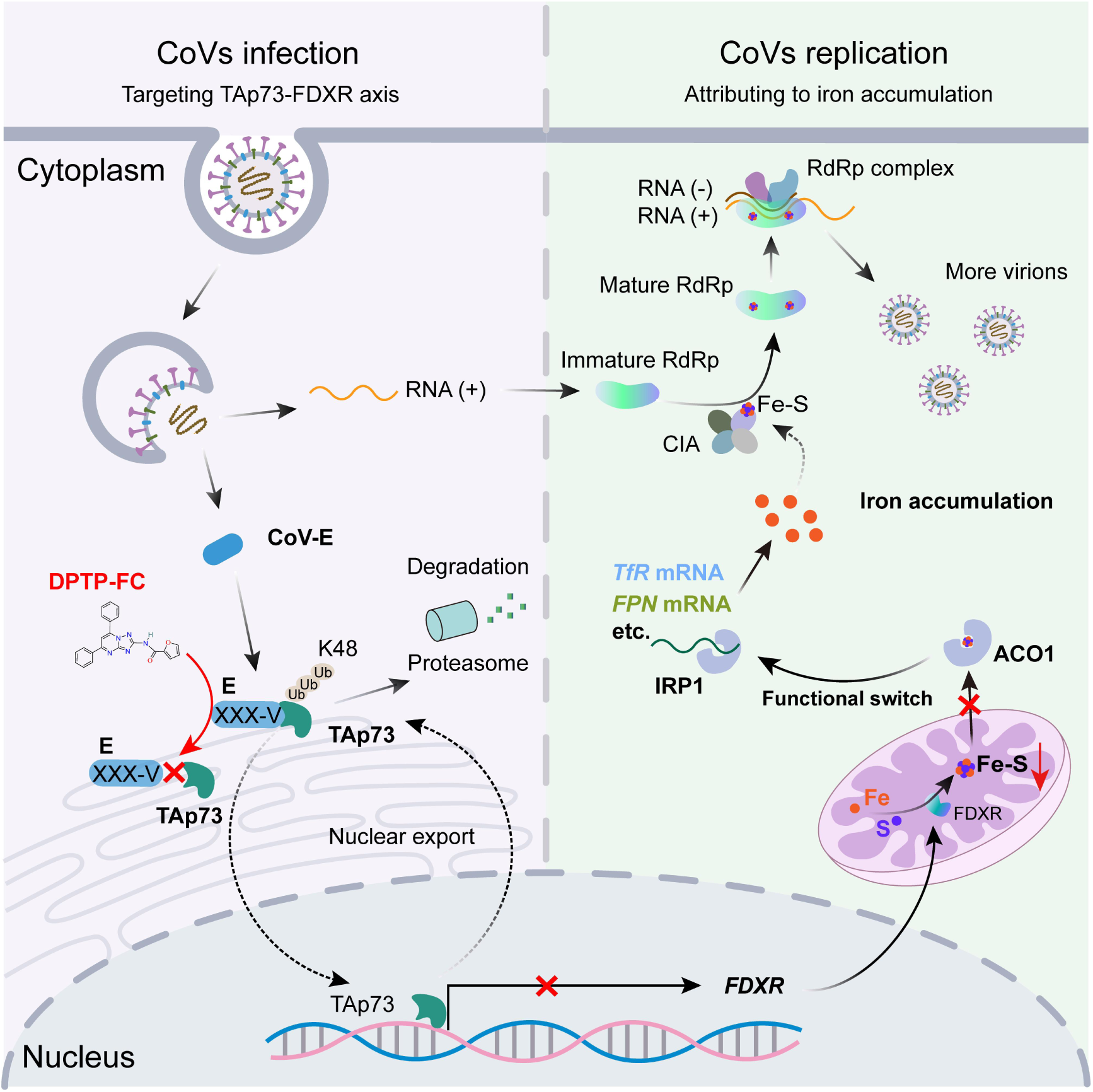
Graphical abstract for CoVs subverts iron homeostasis by hijacking the TAp73-FDXR axis.

This systemic suppression of host Fe-S biogenesis poses a unique challenge for the virus, as the CoVs’ RdRp is strictly dependent on Fe-S clusters for its catalytic integrity^24^. Regarding why FDXR disruption fails to impair Fe-S cluster incorporation into the viral RdRp, despite causing such loss in host proteins, we hypothesize that the intracellular iron-rich environment may facilitate the spontaneous assembly of Fe-S^45^. Alternatively, CoVs might hijack or restructure specific cytoplasmic Fe-S synthesis pathways, thereby acquiring these clusters through direct interaction with the host CIA machinery^46^. This strategy effectively prioritizes the redirection of essential inorganic cofactors toward the viral lifecycle. Nevertheless, whether a specialized, virus-specific Fe-S synthesis machinery is reconstructed within the cytoplasm remains a compelling subject for further investigation.

Our findings identify the CoV-E as the primary upstream orchestrator responsible for establishing this iron-rich environment. Beyond its established roles in virion assembly and ion channel activity^47^, we reveal a previously unrecognized capacity of the CoV-E to manipulate host iron metabolism via TAp73 degradation. Mechanistically, we demonstrate that CoV-E induced nuclear export of TAp73 is a mandatory prerequisite for its proteasomal degradation. While cytoplasmic sequestration might appear a more direct route, we propose that driving active nuclear export confers an evolutionary advantage. It ensures that FDXR transcription is rapidly terminated at the source, thereby accelerating the induction of iron accumulation. This regulatory mechanism is strictly dependent on a conserved valine residue at the C-terminus of the PBM domain, which mediates the K48-linked ubiquitination of TAp73. Notably, the natural divergence of this motif in IBV E leads to a diminished capacity to suppress the TAp73–FDXR axis, indicating that the efficiency of this metabolic hijacking may be a key determinant of virulence across the Coronaviridae family^48^.

Given its central role in metabolic subversion, the E protein represents an attractive target for therapeutic intervention. Traditional antiviral therapies typically target specific viral components^49^, which remains challenging given the high mutation rates of CoVs. In view of metabolic dysregulation can suppress innate immunity, as seen with iron overload^10^, we did indeed observe that CoV, similar to the PEDV, causes an overall suppression of the host’s innate immune pathways (Extended Data Fig. 3a). We therefore propose a metabolic rescue strategy designed to restore host homeostasis and bolster broad-spectrum antiviral immunity. The small molecule DPTP-FC offers a feasible path for restoring iron-sensing fidelity. By binding to the E protein and sterically blocking its interaction with TAp73, DPTP-FC recalibrates the IRP1 sensor and alleviates pathological iron overload. While the resulting antiviral effects likely stem from multifaceted mechanisms, including the alleviation of Iron-induced innate immune suppression, the broad-spectrum potency of DPTP-FC is unequivocal. Given the extensive roles of TAp73 in genomic integrity and programmed cell death^50^, further research is warranted to delineate the long-term physiological consequences of the metabolic rescue strategy on cellular homeostasis. Ultimately, whether other RNA viruses similarly employ sensory deception tactics to hijack constrained host resources remains an intriguing frontier in virus-host coevolution research.

In summary, we demonstrate that the CoV E protein acts as a master regulator that subverts host iron metabolism. These findings define the TAp73–FDXR signaling axis as a critical determinant of viral pathogenesis and identify the restoration of iron-sensing capacity as a promising broad-spectrum antiviral strategy. This metabolic rescue approach represents a departure from traditional antivirals, offering a new avenue to combat current and emerging coronavirus threats.

## Supporting information

Supplementary Data Set

---

**Extended Data Fig. 1:**
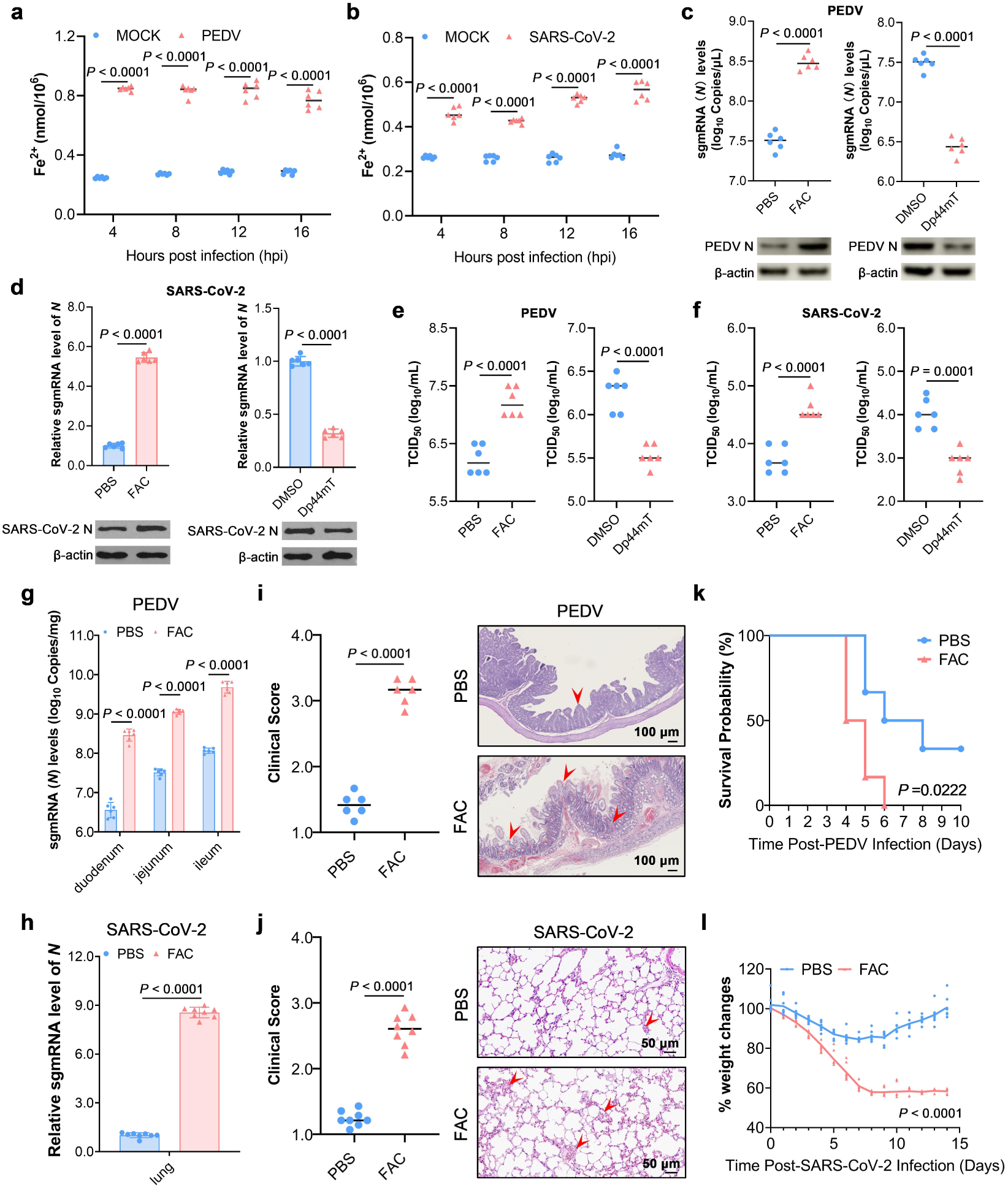
CoVs induce iron overload to promote self-replication. **a, b**, Fe^2+^ levels in PEDV infected IPEC-J2 cells (**a**) and SARS-CoV-2 infected HeLa-ACE2 cells (**b**) for 4 h, 8 h, 12 h and 16 h (*n* = 6). **c**, Absolute quantitative and western blot of PEDV infected IPEC-J2 cells with Dp44mT or FAC treatment for 16 h (*n* = 6). **d**, Relative mRNA levels and western blot analysis of of SARS-CoV-2 infected HeLa-ACE2 cells with Dp44mT or FAC treatment for 16 h (*n* = 6). **e, f**, Virus titer of PEDV infected IPEC-J2 cells (**e**) and SARS-CoV-2 infected HeLa-ACE2 cells (**f**) with Dp44mT or FAC treatment for 16 h (*n* = 6). **g-l**, Viral RNA copy numbers in the small intestines of FAC-handled and control piglets (**g**, *n*=6) and in the lungs of FAC-handled and control hACE2-transgenic mice (**h**, *n*=8); clinical scores with representative haematoxylin and eosin (H&E) staining of jejunal tissues from piglets (**i**, *n*=6) and lung tissues from hACE2-transgenic mice (**j**, *n*=8) showing major pathological lesions indicated by red arrows; survival curves of piglets (**k**, *n*=6) and percentage body weight changes of hACE2-transgenic mice (**l**, *n*=8). Data are represented as mean ± SD, and analyzed by two-tailed Student’s t-test (**a-j, l**) or Gehan-Breslow-Wilcoxon test (**k**).

**Extended Data Fig. 2:**
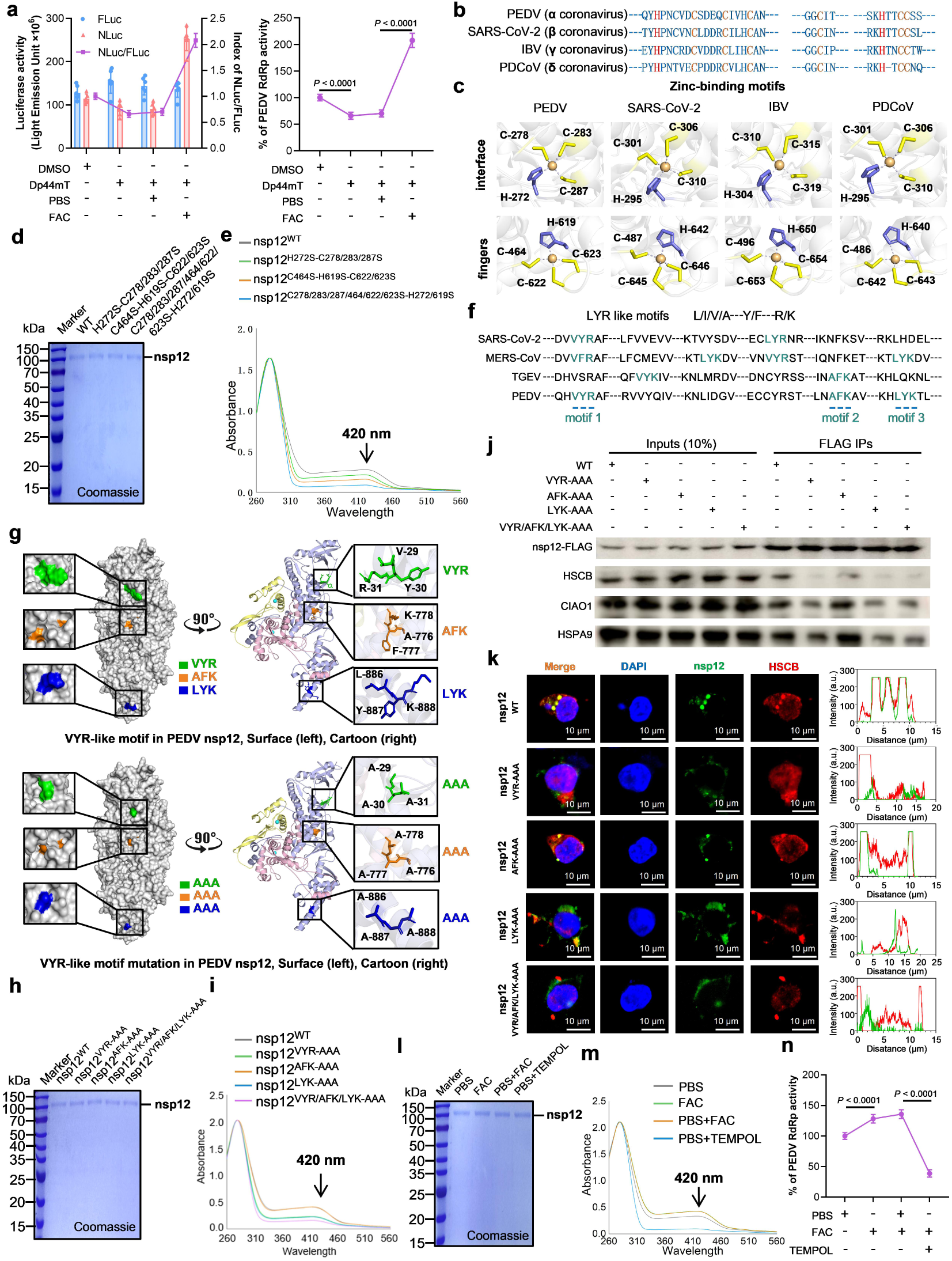
Iron involved in CoVs pathogenicity attributed to the presence of Fe-S in RdRp. **a**, Luciferase activity and PEDV-RdRp activity (NLuc/FLuc) of HEK293T cells treated with Dp44mT or FAC for 16 h (*n*=6). **b**, Sequence alignment of zinc-binding residues in nsp12 from different coronavirus (CoV) genera. **c**, Cartoon representation of homology models of nsp12 highlighting zinc-binding sites. **d-e**, Coomassie Brilliant Blue staining (**d**) and ultraviolet-visible (UV-Vis) absorption spectra **(e**) of anaerobically purified PEDV nsp12 and its cysteine mutants at two CCCH-type zinc-binding motifs. **f**, Sequence alignment of nsp12 from β-coronaviruses (SARS-CoV-2 and MERS-CoV) and α-coronaviruses (TGEV and PEDV), highlighting the conserved LYR motif. **g**, Homology model of PEDV nsp12 showing the spatial arrangement of the LYR-like motif and conformational changes following alanine substitution. **h-i**,Coomassie Brilliant Blue staining (**h**) and UV-Vis absorption spectra (**i**) of anaerobically purified PEDV nsp12 and the indicated LYR-like motif mutants. **j-k**, Immunoprecipitation (IP) and western blot analyses of HSC20, HSPA9, and CIAO1 in IPEC-J2 cells (**j**) and IFA of the interaction and co-localization with HSCB, together with quantification of co-localization (**k**) of PEDV nsp12 or its LYR-like motif mutants. **l-m**, Coomassie Brilliant Blue staining (**l**) and UV-Vis absorption spectra (**m**) of anaerobically purified PEDV nsp12 from PBS, FAC, PBS+FAC, and PBS+TEMPOL-treated samples. **n**, PEDV-RdRp activity of HEK293T cells treated with FAC and TEMPOL for 16 h (*n*=6).Data are represented as mean±SD and analyzed by two-tailed Student’s t-test (**a**, **n**).

**Extended Data Fig. 3:**
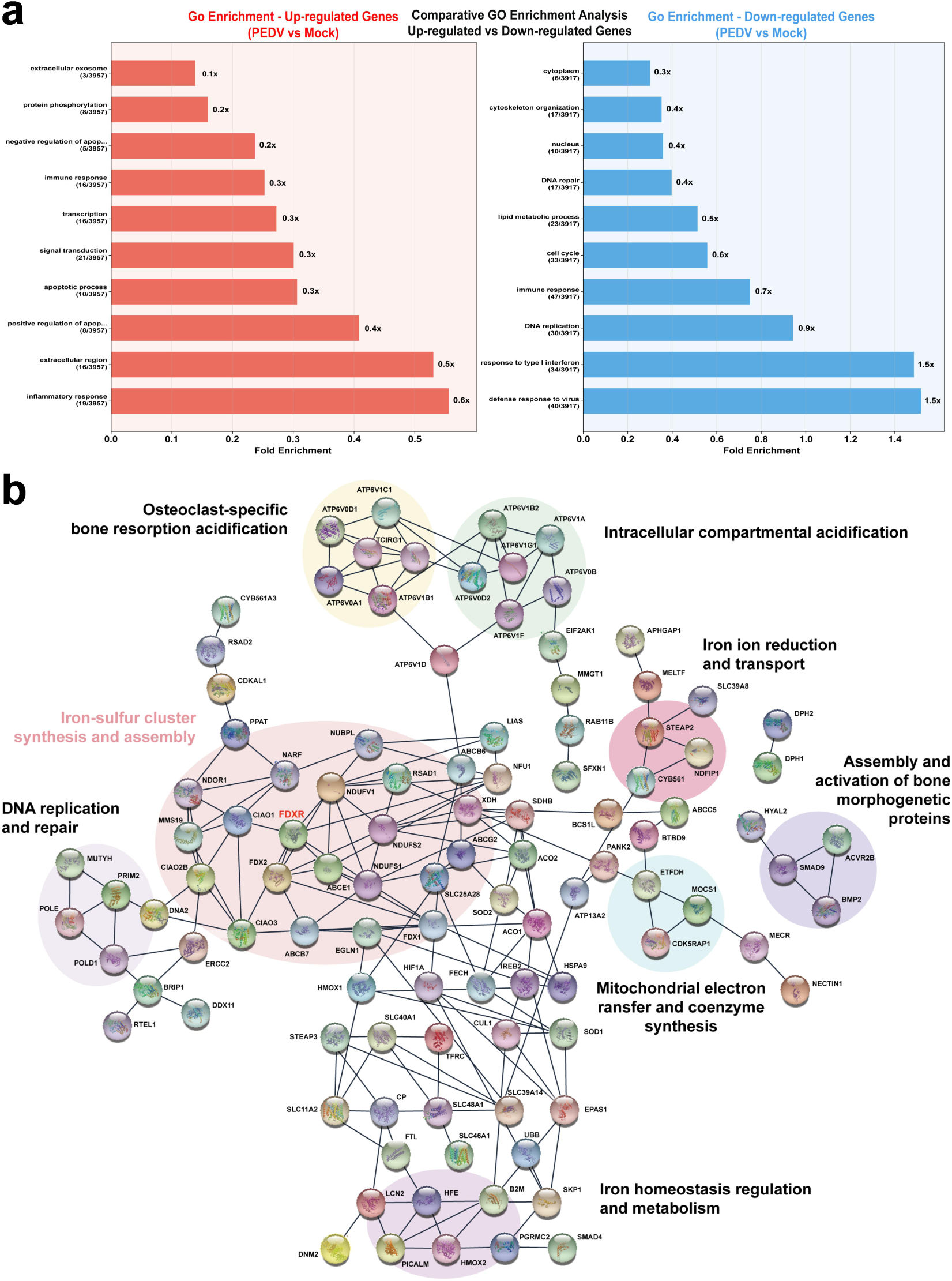
Iron metabolism gene functional enrichment and protein interaction network. **a**. Gene Ontology (GO) enrichment analysis of differentially expressed genes (DEGs). The x-axis represents fold enrichment, and the y-axis represents GO categories. **b**. Protein-protein interaction (PPI) network of iron metabolism-related proteins. Nodes represent proteins, edges indicate known or predicted interactions. FDXR is highlighted in red. Proteins with similar functions have been annotated.

**Extended Data Fig. 4:**
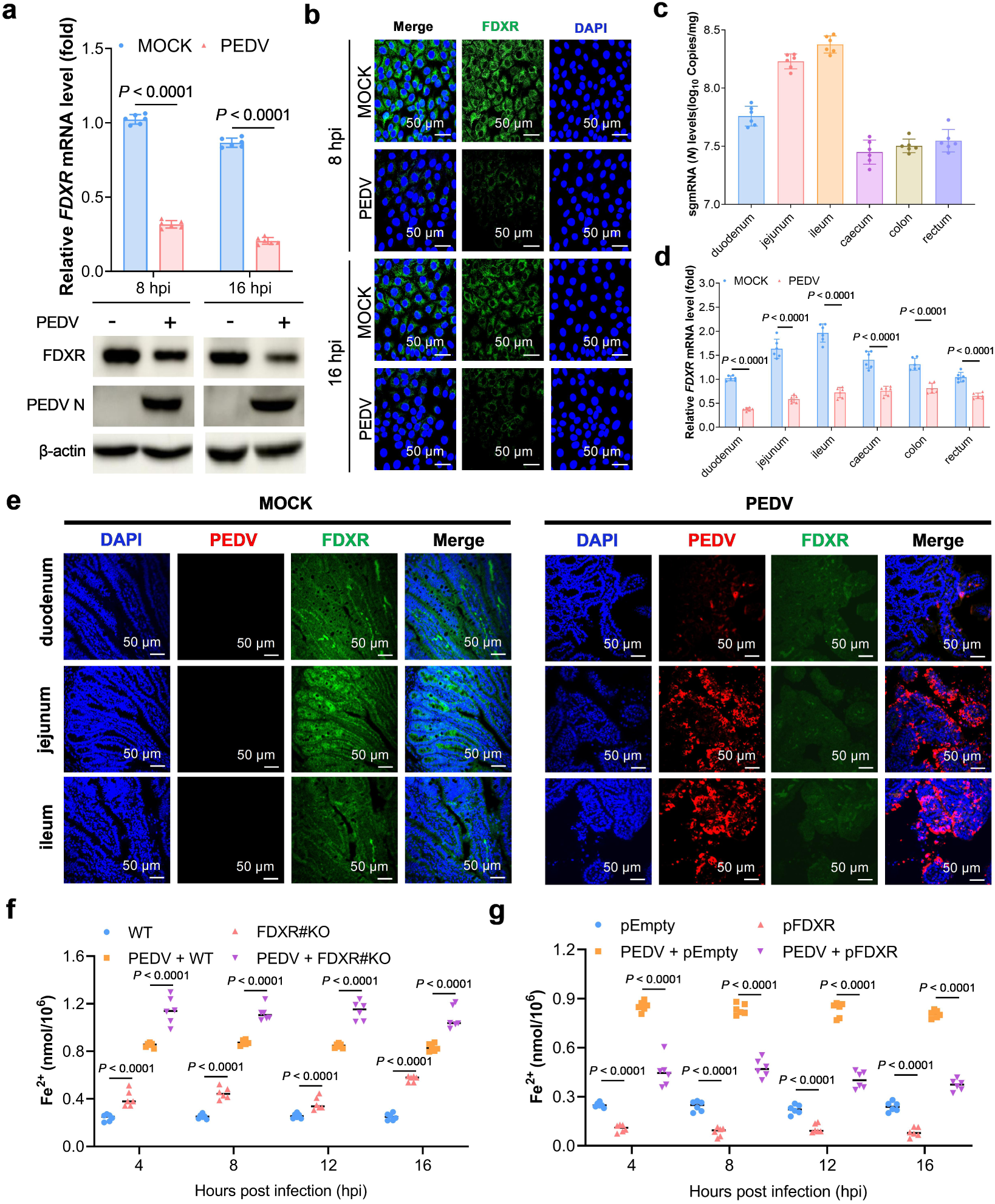
CoVs exploit FDXR to hijack cellular iron for optimal replication. **a**, Relative FDXR mRNA expression and western blot analysis in IPEC-J2 cells infected with PEDV strain LZ202401 (MOI 1) at 8 and 16 h post infection. **b**, IFA analysis of IPEC-J2 cells infected with PEDV strain LZ202401 (MOI 1) at 8 and 16 h, showing PEDV N protein (red) and FDXR (green)(*n* = 6). **c-e**, Analyses of PEDV strain LZ202401 infection in piglets at 7 dpi, including absolute quantification of PEDV N gene copy numbers per milligram of tissue (**c**), relative quantification of *FDXR* expression in tissues (**d**), and IFA of small intestinal tissues showing PEDV N protein (red) and FDXR (green) (**e**, *n* = 6). **f**, Fe^2+^ levels in IPEC-J2 cells and FDXR knockout (FDXR#KO) IPEC-J2 cells with or without PEDV infection for 4 h, 8 h, 12 h and 16 h (*n* = 6). **g**, Fe^2+^ levels in pEmpty or pFDXR -transfected IPEC-J2 cells with or without PEDV infection for 4 h, 8 h, 12 h and 16 h (*n* = 6). Data are represented as mean ± SD, and analyzed by two-tailed Student’s t-test (**a, d, f, g**).

**Extended Data Fig. 5:**
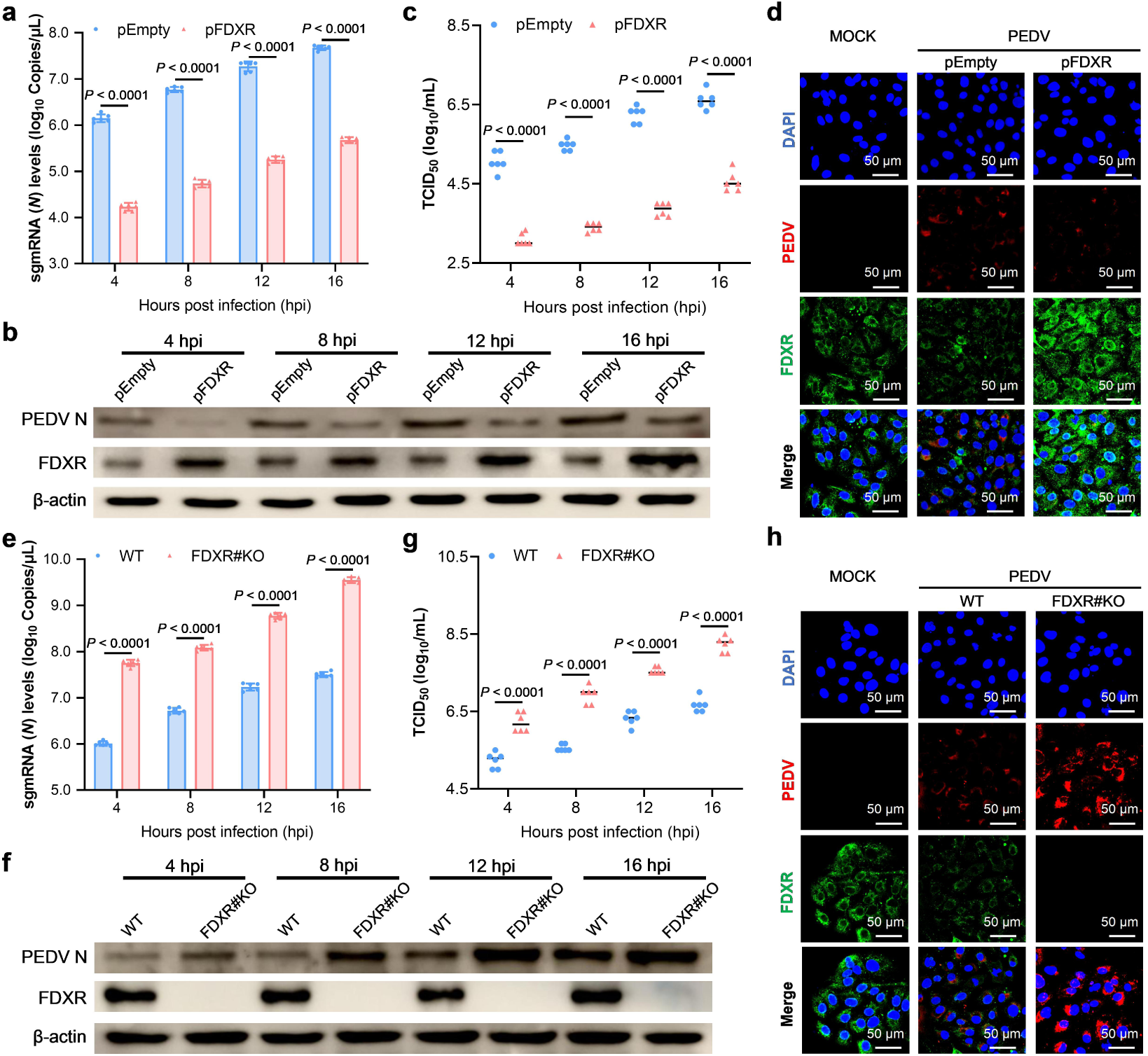
FDXR suppresses PEDV replication. **a-d**, Effects of FDXR overexpression on PEDV replication in IPEC-J2 cells. Cells transfected with pEmpty or pFDXR were infected or mock-infected with PEDV and analysed at 4, 8, 12 and 16 h. PEDV N gene copy numbers were quantified by qRT-PCR (**a**), N protein expression was analysed by western blotting (**b**), viral titres were determined by TCID₅₀ assay (**c**), and IFA showed PEDV N protein (red) and FDXR (green) (**d**, *n* = 6). **e-h**, Effects of FDXR knockout on PEDV replication in IPEC-J2 cells. Wild-type and FDXR#KO IPEC-J2 cells were infected or mock-infected with PEDV and analysed at 4, 8, 12 and 16 h. PEDV N gene copy numbers were quantified by qRT-PCR (**e**), N protein expression was analysed by western blotting (**f**), viral titres were determined by TCID₅₀ assay (**g**), and IFA showed PEDV N protein (red) and FDXR (green) (**h**, *n* = 6). Data are represented as mean ± SD, and analyzed by two-tailed Student’s t-test (**a, c, e, g**).

**Extended Data Fig. 6:**
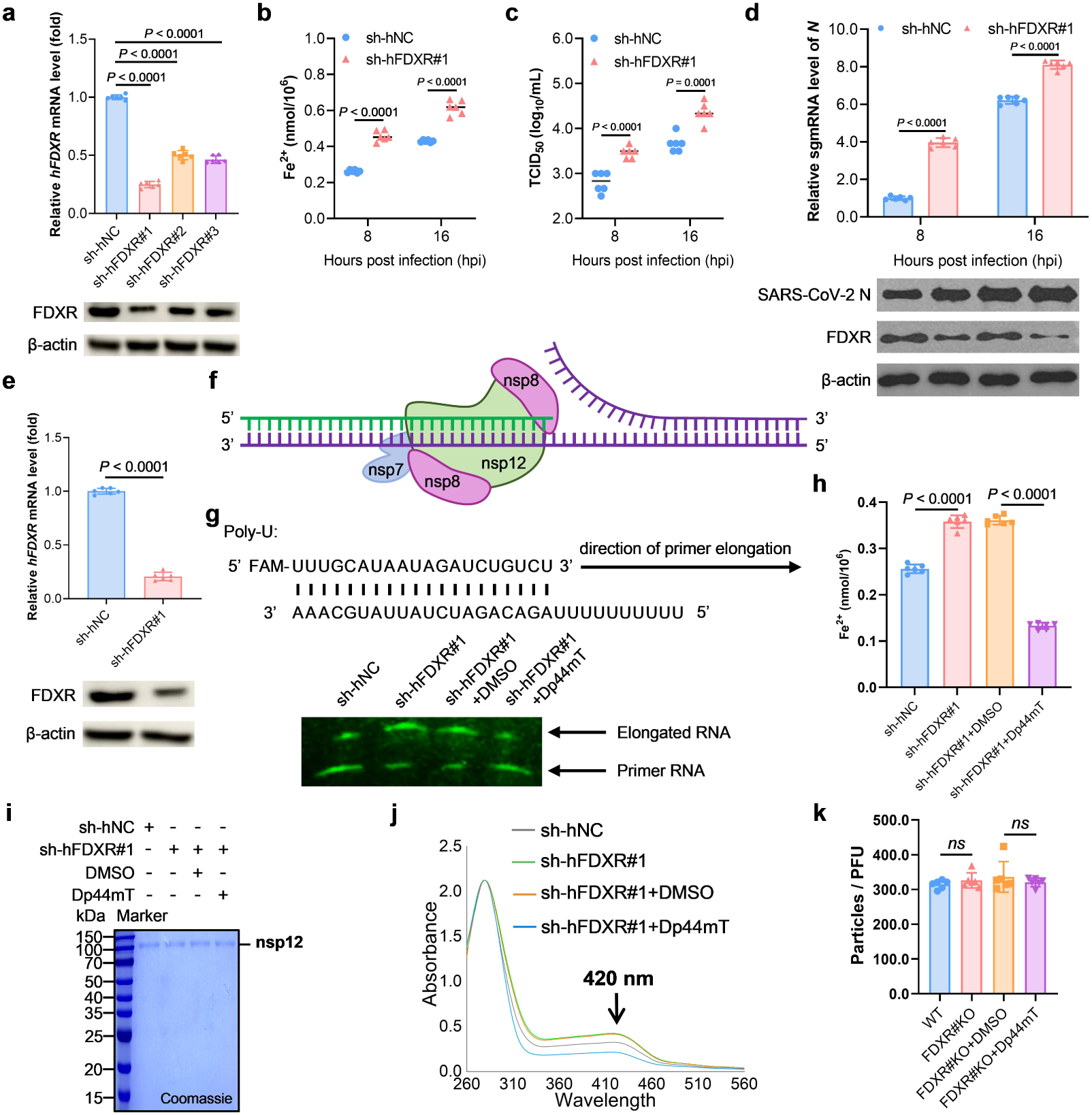
FDXR regulates the synthesis of Fe-S in SARS-CoV-2-RdRp via modulating cellular iron. **a**, Relative levels of *hFDXR* in HeLa-ACE2 cells with sh-hNC or sh-hFDXR transfection for 16 h (*n* = 6). **b-d**, Fe^2+^ levels (**b**), virus titer (**c**), Relative mRNA levels and western blot analysis of (**d**) analysis of SARS-CoV-2 infected HeLa-ACE2 cells with sh-hNC or sh-hFDXR#1 transfection for 8 h and 16 h (*n* = 6). **e**, Relative levels of *hFDXR* in Expi293F cells with sh-hNC or sh-hFDXR#1 transfection for 16 h (*n* = 6). **f, g**, Elongation of the partial RNA duplex by the purified RdRp complex treated with shFDXR#1 alone or in combination with Dp44mT. **h**, Fe^2+^ levels of sh-hNC- or sh-hFDXR#1-transfected Expi293F cells in the presence or absence of Dp44mT for 16 h (*n* = 6). **i, j**, Coomassie Brilliant Blue staining (**i**) and UV-Vis absorption spectra (**j**) of anaerobically purified ACO1/IRP1 in sh-hNC- or sh-hFDXR#1-transfected Expi293F cells in the presence or absence of Dp44mT. **k**, Plaque formation and transmission electron microscopy analysis of the number of PEDV viral particles in IPEC-J2 cells and FDXR#KO IPEC-J2 cells in the presence or absence of Dp44mT (*n* = 6). The proportions of defective virus particles were expressed as the percentage of infectious virus particles in the total number of virus particles. Data are represented as mean ± SD, and analyzed by one-way ANOVA with Tukey’s multiple comparisons test (**a**) or two-tailed Student’s t-test (**b-e, h, k**).

**Extended Data Fig. 7:**
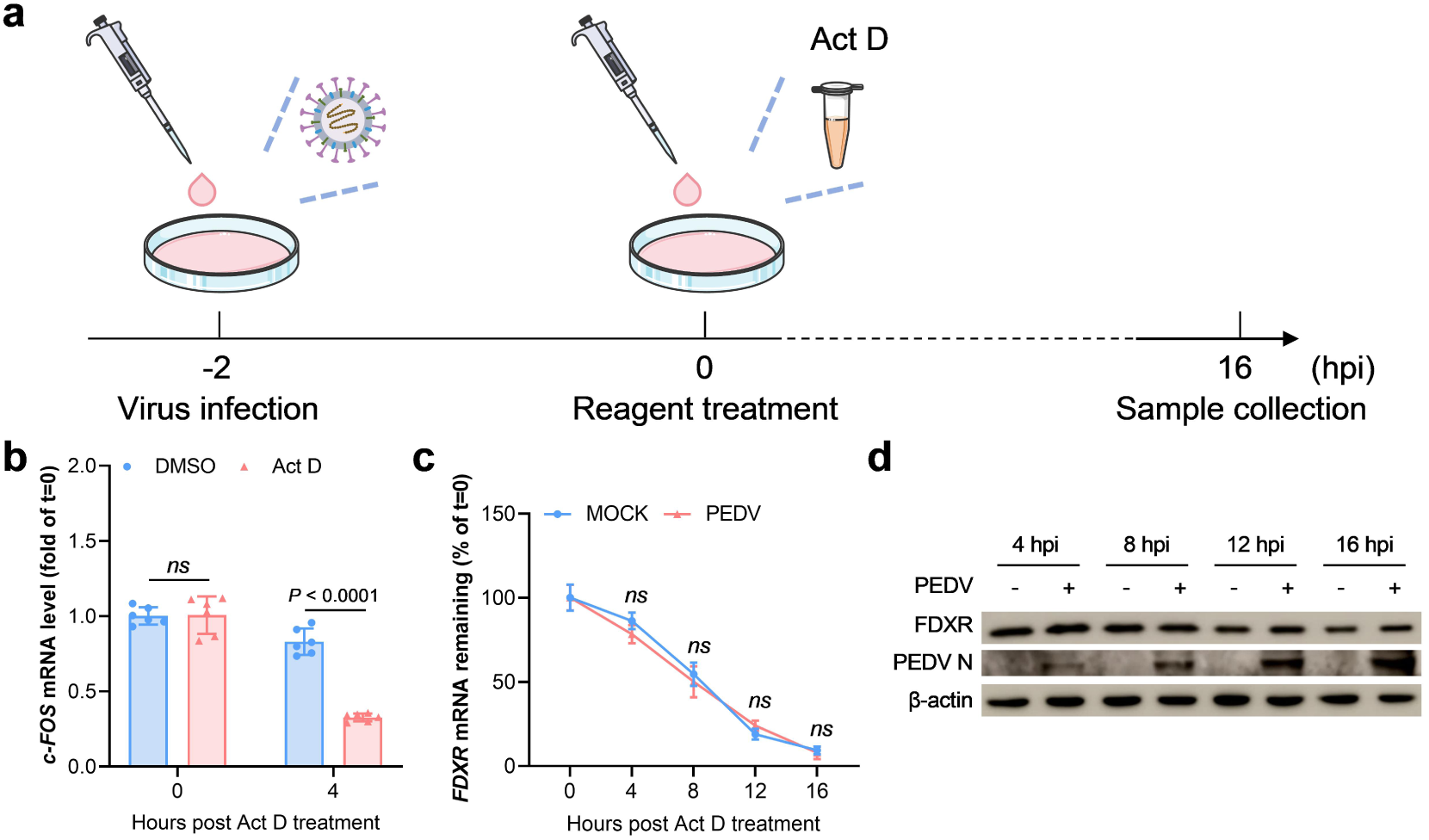
*FDXR* mRNA exhibits high stability under transcriptional blockade. **a**, Experimental setup for Act D treament. **b-d**, *c-fos* (**b**), *FDXR* (**c**) levels and western blot (**d**) analysis of analysis of IPEC-J2 cells with Act D treament for the specified time (*n* = 6). Data are represented as mean ± SD, and analyzed by two-tailed Student’s t-test (**b, c**).

**Extended Data Fig. 8:**
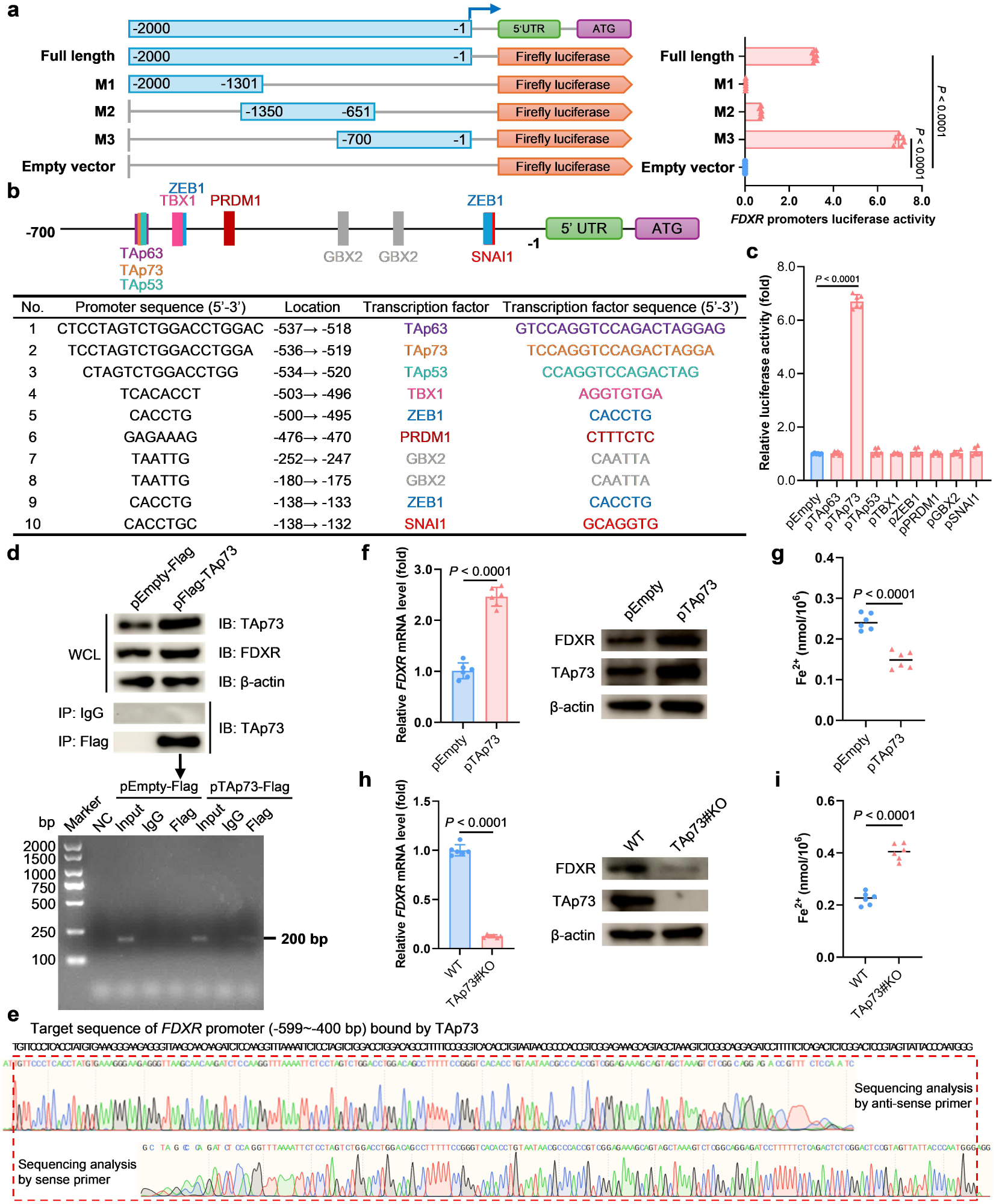
TAp73 drives *FDXR* promoter activity. **a**, Schematic diagram of full-length and truncated *FDXR* promoters (designated M1-M3) and its luciferase activity in HEK293T cells for 16 h (*n* = 6). **b**, Potential transcription factors on *FDXR* promoter predicted by the JASPAR database. **c**, Relative luciferase activity of *FDXR* promoters (M3) in HEK293T cells with overexpressed transcription factors for 16 h (*n* = 6). **d**, ChIP analysis of the binding of TAp73 protein and *FDXR* mRNA in IPEC-J2 cells. **e**, Sequence of ChIP products derived from pTAp73-Flag. **f-i**, Relative levels, western blot (**f, h**) and Fe^2+^ levels (**g, i**) analysis of IPEC-J2 cells with TAp73 overexpressed or KO for 16 h (*n* = 6). Data are represented as mean ± SD, and analyzed by one-way ANOVA with Tukey’s multiple comparisons test (**a, c**) or two-tailed Student’s t-test (**f-i**).

**Extended Data Fig. 9:**
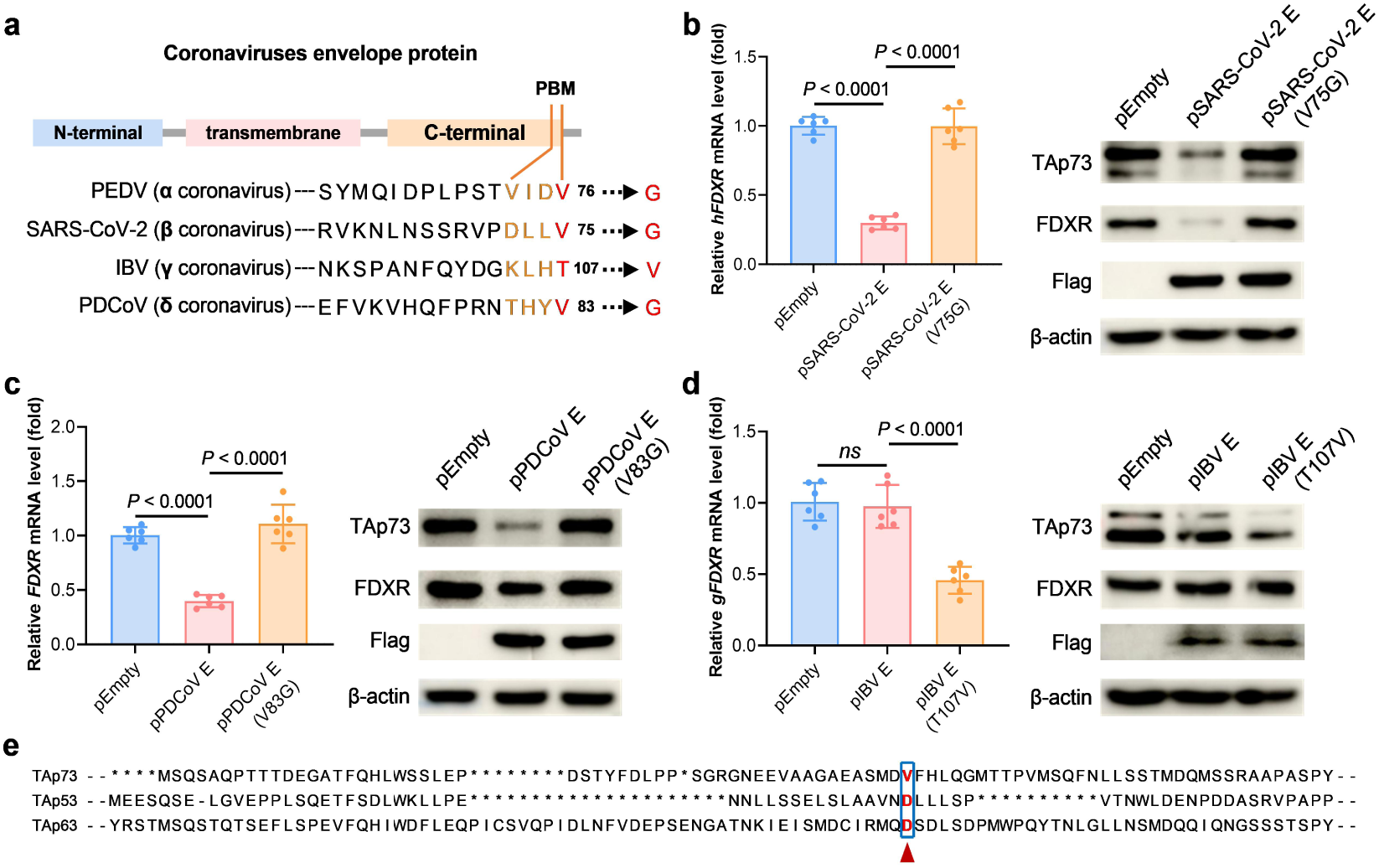
CoVs E mediate TAp73 degradation through valine at the C-terminal of its PBM domain. **a**, Domain analysis of CoVs E from each genus. **b**, Relative mRNA levels and western blot analysis of analysis of HeLa-ACE2 cells with SARS-CoV-2 E or SARS-CoV-2 E (V75G) transfection for 16 h (*n* = 6). **c**, Relative mRNA levels and western blot analysis of analysis of LLC-PK1 cells with PDCoV E or PDCoV E (V83G) transfection for 16 h (*n* = 6). **d**, Relative mRNA levels and western blot analysis of analysis of DF1 cells with IBV E or IBV E (T107V) transfection for 16 h (*n* = 6). **e**, Amino acid sequence alignment of TAp53, TAp63 and TAp73, highlighting the non-conserved motifs interacting with PEDV E protein.. Data are represented as mean ± SD, and analyzed by one-way ANOVA with Tukey’s multiple comparisons test (**b-d**).

**Extended Data Fig. 10:**
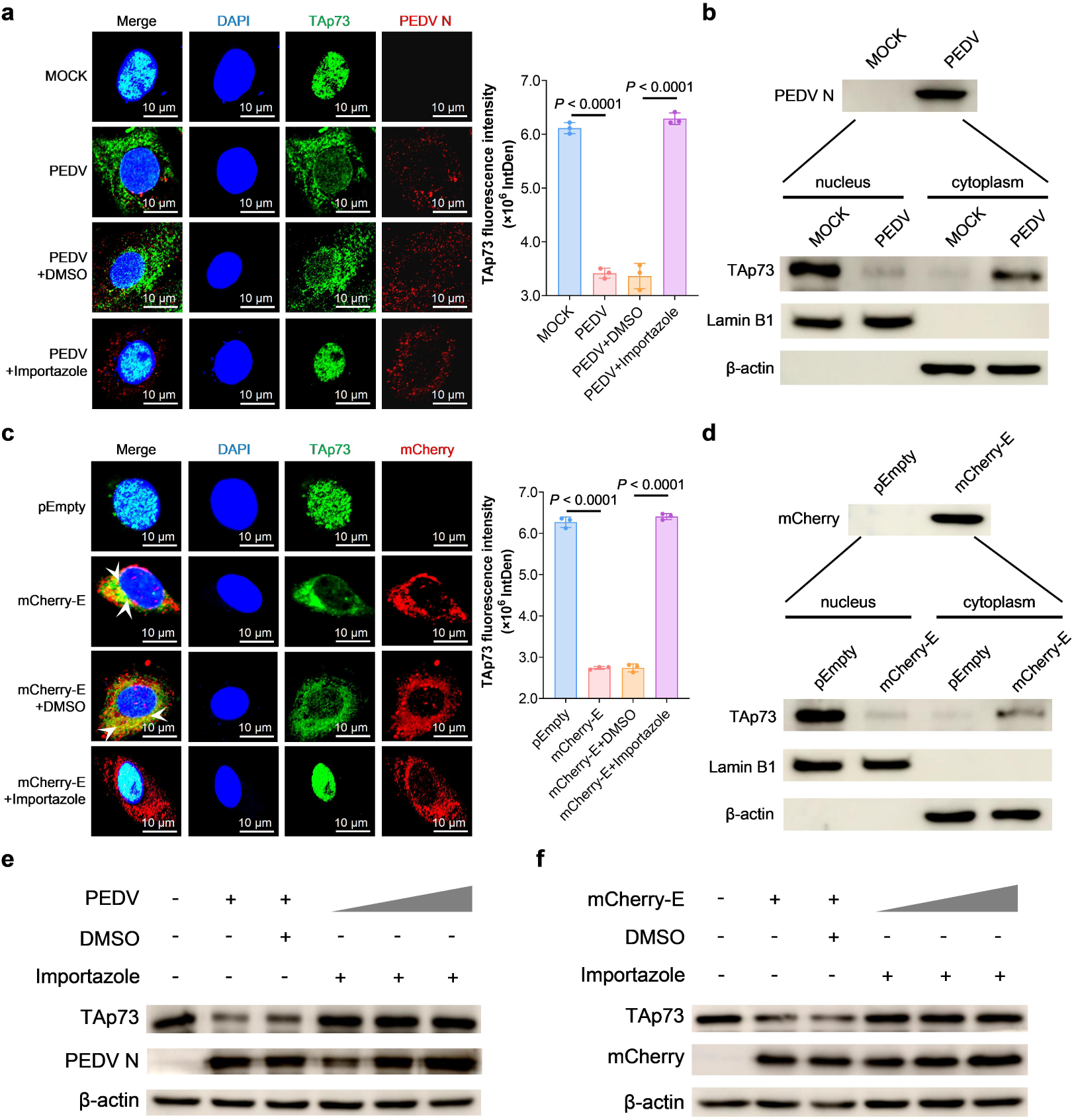
PEDV E protein induces TAp73 nuclear translocation. **a**, IFA analysis of TAp73 nuclear export in IPEC-J2 cells under MOCK, PEDV, PEDV+DMSO, and PEDV+importazole treatment conditions. TAp73 is shown in green and PEDV N protein in red (left). Quantification of nuclear TAp73 fluorescence intensity is shown on the right (*n* = 6). **b**, Western blot analysis of TAp73 protein levels in nuclear and cytoplasmic fractions of cells following PEDV infection. Lamin B1 and β-actin were used as nuclear and cytoplasmic loading controls, respectively(*n* = 6). **c**, IFA analysis of TAp73 nuclear export in cells transfected with pEmpty, mCherry-E, mCherry-E+DMSO, or mCherry-E+importazole. TAp73 is shown in green and mCherry-E in red (left). Quantification of nuclear TAp73 fluorescence intensity is shown on the right. **d**, Western blot analysis of TAp73 protein levels in nuclear and cytoplasmic fractions of cells transfected with mCherry-E. Lamin B1 and β-actin were used as nuclear and cytoplasmic loading controls, respectively. **e, f**, Western blot analysis of IPEC-J2 cells co-treated with importazole and either dose-dependent PEDV infection (**e**) or PEDV E transfection (**f**) for 16 h. Data are represented as mean ± SD, and analyzed by two-tailed Student’s t-test (**a, c**).

**Extended Data Fig. 11:**
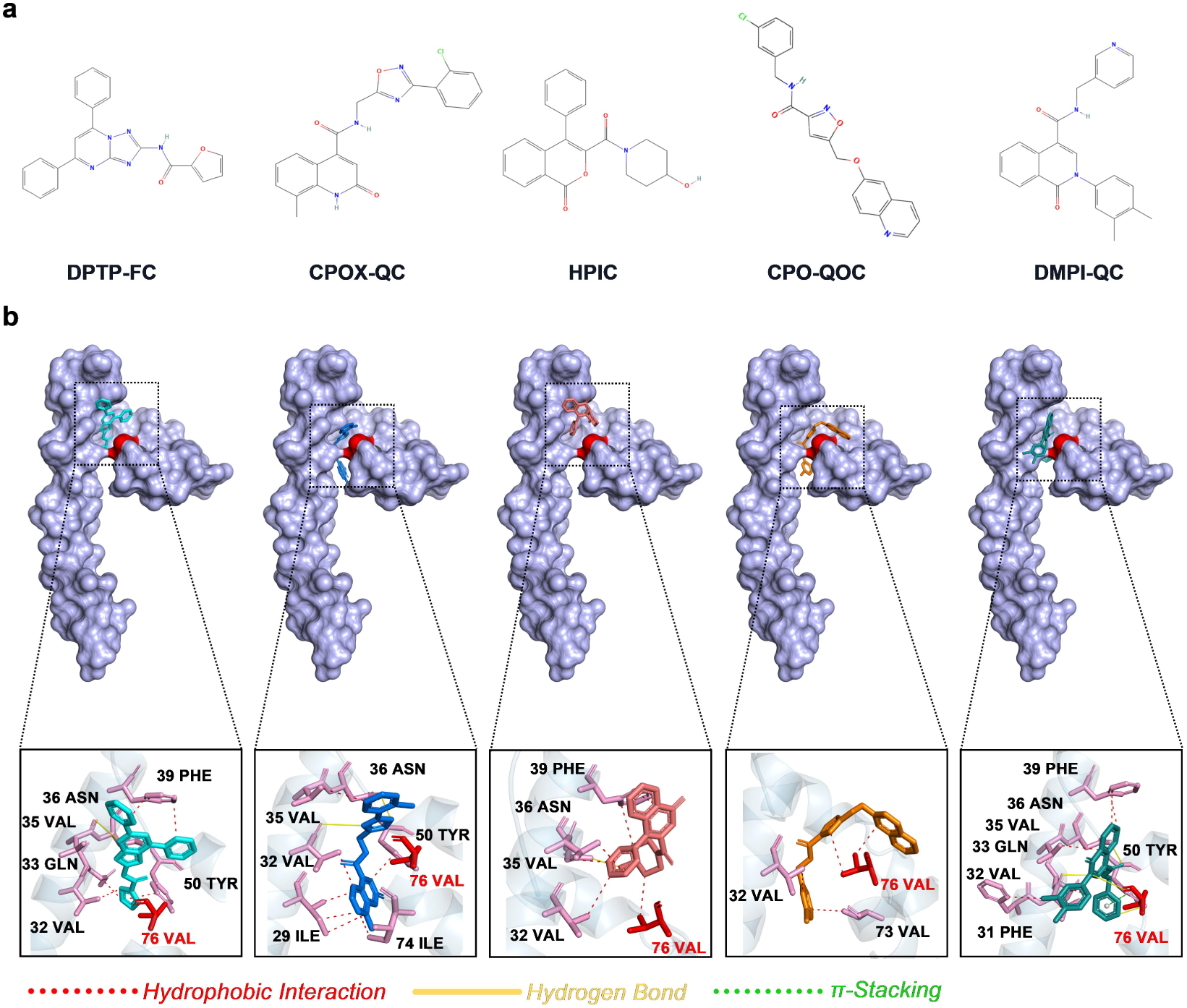
Structure-based docking of small-molecule compounds targeting V76 of PEDV E protein. **a**, Chemical structures of the top five small-molecule compounds with the lowest docking energies targeting V76 of the PEDV E protein. **b**, Docking visualizations of the optimal binding conformations of the top five small-molecule compounds with the lowest docking energies targeting V76 (red), of the PEDV E protein. Hydrophobic interactions are indicated by red dashed lines, hydrogen bonds by yellow solid lines, and π-stacking interactions by green dashed lines; all intermolecular interactions between the compounds and the PEDV E protein are shown.

**Extended Data Fig. 12:**
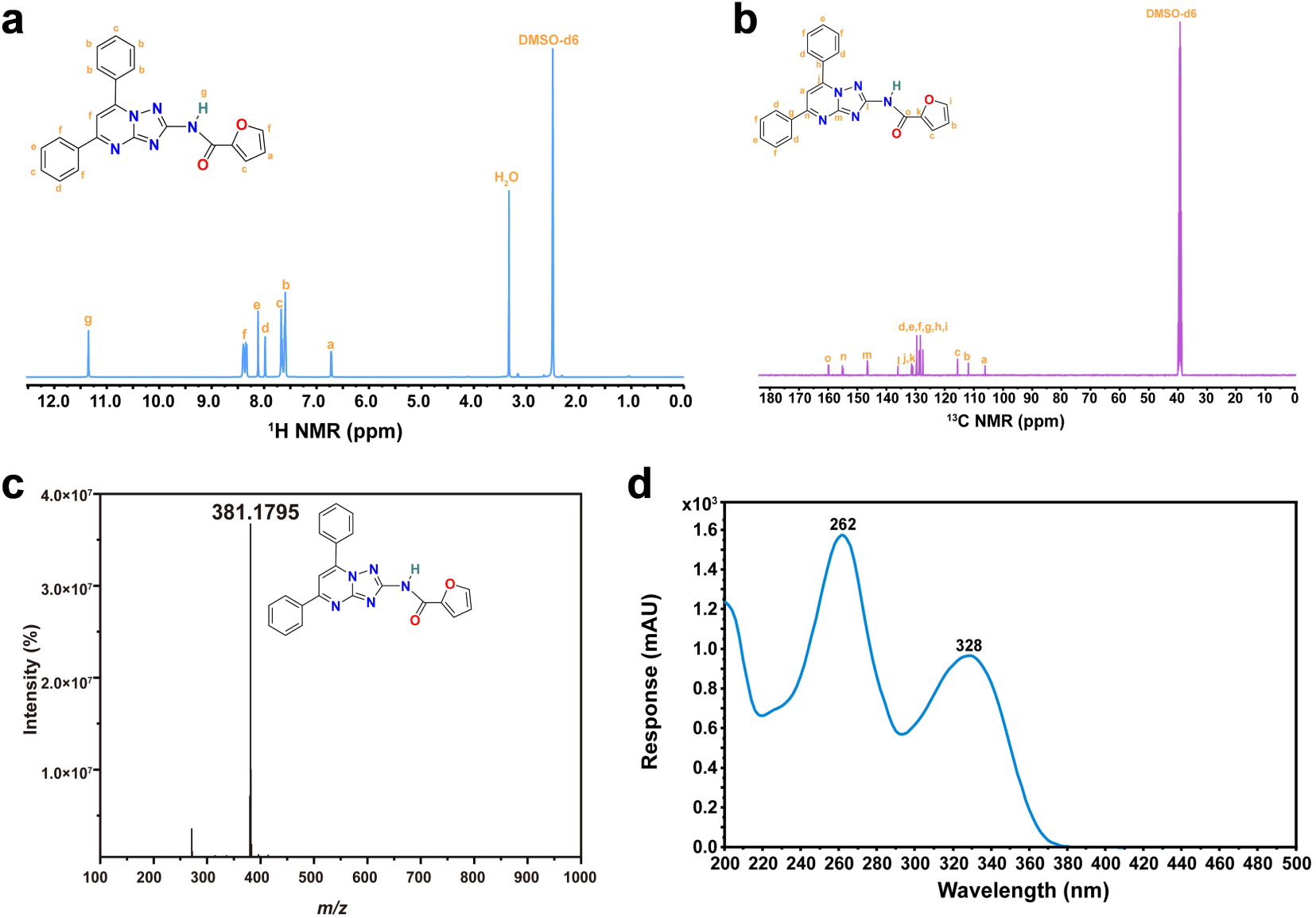
The structural characterization of DPTP-FC. **a-d,** ^1^H NMR (**a**) and ^13^C NMR (**b**) analysis of DPTP-FC in DMSO-d_6_. HRMS (**c**) and ultraviolet absorption spectra (**d**) of DPTP-FC.

**Extended Data Fig. 13:**
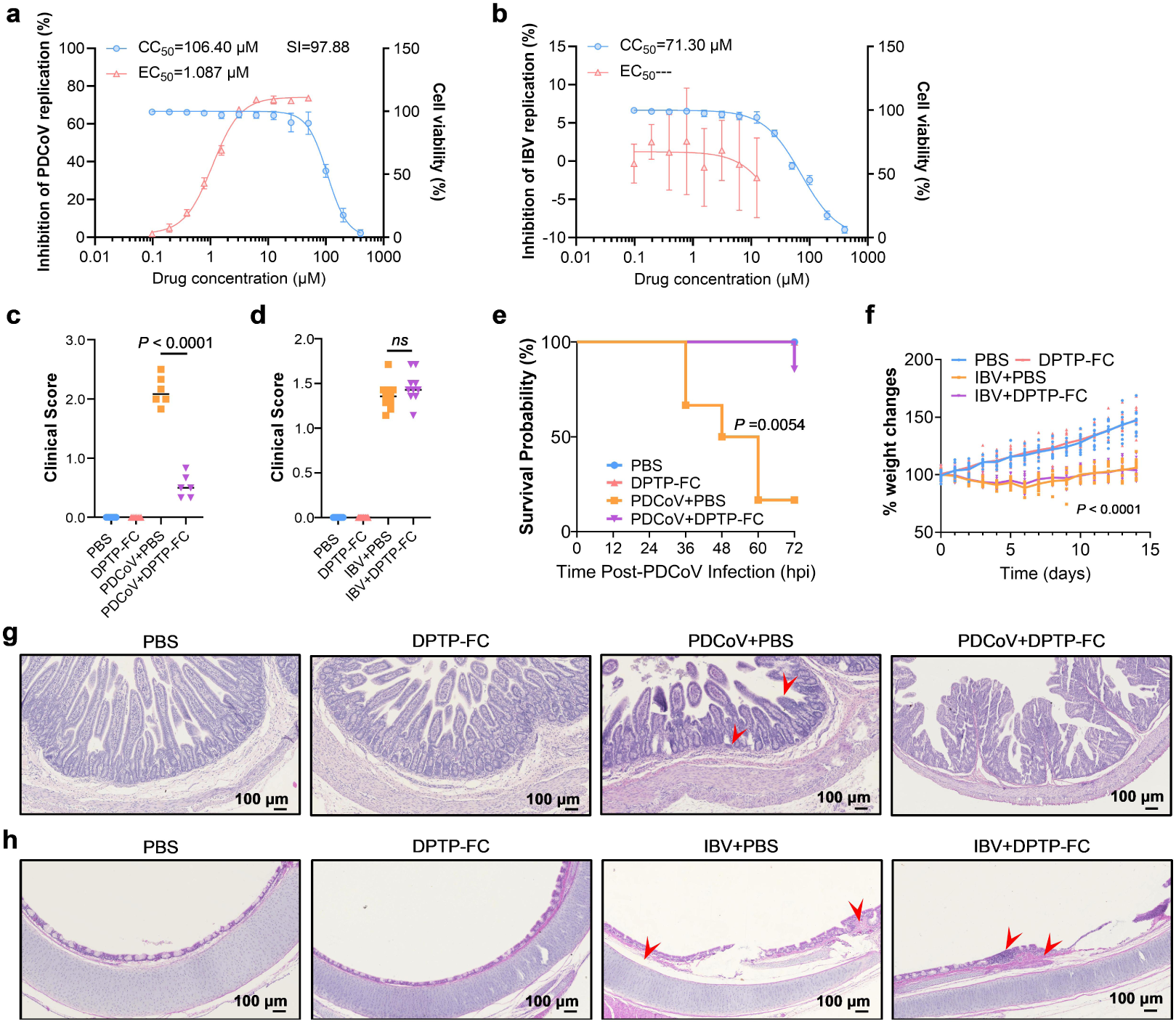
In vivo anti-IBV and anti-PDCoV activity of DPTC-FC. **a-h**, Antiviral efficacy of DPTP-FC against PDCoV and IBV in vitro and in vivo.The EC_50_ of DPTP-FC against PDCoV and the CC_50_, as determined in LLC-PK1 cells (**a**), and the EC_50_ and CC_50_ values of DPTP-FC against IBV, as measured in DF1 cells (**b**). Clinical scores of PDCoV-infected piglets with or without DPTP-FC treatment (**c**, *n*=6) and IBV-infected chickens with or without DPTP-FC treatment (**d**, *n*=10). Survival curves of PDCoV-infected piglets with or without DPTP-FC treatment (**e**, *n*=6) and percentage body weight changes of IBV-infected chickens with or without DPTP-FC treatment (**f**, *n*=10). H&E staining of jejunal tissues from PDCoV-infected piglets treated with or without DPTP-FC (**g**) and tracheal tissues from IBV-infected chickens (**h**), with representative pathological lesions indicated by red arrows.Data are presented as mean±SD and were analysed by two-tailed Student’s t-test (**c, d, f**) or the Gehan-Breslow-Wilcoxon test (**e**).

**Extended Data Table 1:** Docking-based ranking and interaction analysis of small-molecule compounds targeting V76 of PEDV E protein.

| Compound | Energy | nRot | HBA | HBD | LogP | MW | TPSA |
| --- | --- | --- | --- | --- | --- | --- | --- |
| 860538_Intermediate | -8.3 | 4 | 7 | 1 | 3.99 | 381.3868 | 91.64 |
| 26647294_Intermediate | -8 | 4 | 7 | 2 | 3.85 | 394.8111 | 100.88 |
| 7965023_Accepted | -7.9 | 2 | 5 | 1 | 2.63 | 349.37986 | 70.75 |
| 24799473_Intermediate | -7.8 | 6 | 6 | 1 | 3.87 | 393.82304 | 77.25 |
| 24360339_Intermediate | -7.8 | 4 | 5 | 1 | 3.22 | 383.44244 | 63.99 |
| 17402129_Accepted | -7.8 | 2 | 4 | 1 | 3.43 | 319.3323632 | 58.11 |
| 26539518_Intermediate | -7.7 | 3 | 6 | 1 | 2.5 | 370.35756 | 79.62 |
| 17450998_Accepted | -7.7 | 4 | 7 | 1 | 4 | 395.41338 | 88.55 |
| 17402945_Intermediate | -7.6 | 2 | 6 | 0 | 2.28 | 355.34622 | 77.99 |
| 14739966_Intermediate | -7.6 | 0 | 6 | 2 | 2.9 | 429.51088 | 96.42 |
| 26649115_Accepted | -7.6 | 3 | 6 | 1 | 1.83 | 392.49402 | 62.66 |
| 24801000_Intermediate | -7.6 | 5 | 6 | 1 | 3.75 | 397.3796 | 94.58 |
| 26728213_Intermediate | -7.6 | 2 | 6 | 1 | 3.95 | 383.39938 | 75.44 |

| Hydrophobic Interaction |  |  |  | Hydrogen Bond |  |  |  |  |  |
| --- | --- | --- | --- | --- | --- | --- | --- | --- | --- |
| Index | Residue | AA | Distance | Index | Residue | AA | Distance H-A | Distance D-A | Donor Angle |
| 1 | 32 | VAL | 3.62 | 1 | 35 | VAL | 3.71 | 4.06 | 103.98 |
| 1 | 33 | GLN | 3.66 | 1 | 36 | ASN | 1.96 | 2.9 | 157.82 |
| 1 | 39 | PHE | 3.59 |  |  |  |  |  |  |
| 1 | 39 | PHE | 3.47 |  |  |  |  |  |  |
| 1 | 50 | TYR | 3.86 |  |  |  |  |  |  |
| Index | Residue | AA | Distance | Index | Residue | AA | Distance H-A | Distance D-A | Donor Angle |
| 2 | 29 | ILE | 3.87 | 2 | 32 | VAL | 2.78 | 3.57 | 137.27 |
| 2 | 29 | ILE | 3.58 | 2 | 36 | ASN | 2.39 | 3.01 | 120.38 |
| 2 | 29 | ILE | 3.69 |  |  |  |  |  |  |
| 2 | 32 | VAL | 3.64 |  |  |  |  |  |  |
| 2 | 35 | VAL | 3.83 |  |  |  |  |  |  |
| 2 | 50 | TYR | 3.62 |  |  |  |  |  |  |
| 2 | 74 | ILE | 3.35 |  |  |  |  |  |  |
| Index | Residue | AA | Distance | Index | Residue | AA | Distance H-A | Distance D-A | Donor Angle |
| 3 | 32 | VAL | 3.91 | 3 | 36 | ASN | 2.47 | 3.12 | 123.49 |
| 3 | 35 | VAL | 3.66 |  |  |  |  |  |  |
| 3 | 36 | ASN | 3.75 |  |  |  |  |  |  |
| 3 | 39 | PHE | 3.77 |  |  |  |  |  |  |
| 3 | 39 | PHE | 3.56 |  |  |  |  |  |  |
| 3 | 76 | VAL | 3.69 |  |  |  |  |  |  |

| Hydrophobic Interaction |  |  |  |
| --- | --- | --- | --- |
| Index | Residue | AA | Distance |
| 4 | 32 | VAL | 3.47 |
| 4 | 73 | VAL | 3.69 |
| 4 | 76 | VAL | 3.35 |
| 4 | 76 | VAL | 3.75 |

| Hydrophobic Interaction |  |  |  | Hydrogen Bond |  |  |  |  |  |
| --- | --- | --- | --- | --- | --- | --- | --- | --- | --- |
| Index | Residue | AA | Distance | Index | Residue | AA | Distance H-A | Distance D-A | Donor Angle |
| 5 | 31 | PHE | 3.75 | 5 | 32 | VAL | 3.13 | 4.03 | 148.8 |
| 5 | 32 | VAL | 3.68 | 5 | 36 | ASN | 2.53 | 3.22 | 127.07 |
| 5 | 33 | GLN | 3.65 | 5 | 50 | TYR | 3.16 | 3.8 | 127.42 |
| 5 | 35 | VAL | 3.86 | 5 | 76 | VAL | 2.84 | 3.48 | 122.74 |
| 5 | 39 | PHE | 3.71 |  |  |  |  |  |  |
| 5 | 39 | PHE | 3.77 |  |  |  |  |  |  |
| π-Stacking | | | | | | | | | |
| Index | Residue | AA | Distance | Angle | Offset |  |  |  |  |
| 5 | 50 | TYR | 4.92 | 72.81 | 1.07 |  |  |  |  |

## Methods

### Cell lines and culture conditions

Porcine intestinal epithelial cells (IPEC-J2), FDXR knockout IPEC-J2 cells (FDXR#KO), and TAp73 knockout IPEC-J2 cells (TAp73#KO) were maintained in Dulbecco’s modified Eagle medium/Nutrient Mixture F-12 (DMEM/F-12; Pricella, PM150312) supplemented with 10% fetal bovine serum (FBS; Biological Industries) and 100 U/mL penicillin-streptomycin. HeLa cells stably expressing human ACE2 (HeLa-ACE2) were generated by lentiviral transduction in a biosafety level-3 (BSL-3) facility at Shenzhen Third People’s Hospital. Vero (ATCC CCL-81), VeroE6 (ATCC CRL-1586), HEK293T (ATCC CRL-3216), Expi293F, LLC-PK1, and DF-1 cells were cultured in DMEM (Pricella, PM150210) supplemented with 10% FBS and antibiotics unless otherwise indicated. HEK293F cells were maintained in SMM 293-TII complete medium (Sino Biological, M293TII). All cell lines were cultured at 37 ℃ in a humidified incubator with 5% CO₂ and were routinely tested and confirmed negative for mycoplasma contamination.

### Viruses

PEDV LZ202401 (GenBank: PP999774) was propagated in Vero cells in the presence of 10 µg/mL trypsin. Viral titres were determined by plaque assay. SARS-CoV-2 strain SZTH-003 (GISAID: EPI_ISL_406594) was isolated from a COVID-19 patient and amplified once in VeroE6 cells under BSL-3 conditions. Infectious bronchitis virus (IBV) SX2024 (GenBank: PV185508) was propagated in 10-day-old SPF embryonated chicken eggs. Porcine deltacoronavirus (PDCoV) CZ2020 was amplified in LLC-PK1 cells. Viral titres were determined by plaque assay, TCID_50_ or EID_50_ as indicated^51,52^.

### Animals

All animal experiments were approved by the institutional animal care and use committees and conducted in accordance with national guidelines. Three-day-old piglets (Sus scrofa domesticus) were obtained from a commercial SPF farm in Shaanxi Province, China. Piglets were housed in temperature- and humidity-controlled pathogen-free facilities with ad libitum access to milk replacer and water. Six-day-old K18-hACE2 transgenic mice were purchased from Cyagen Biosciences and maintained under SPF conditions. Fertilized White Leghorn chicken eggs were obtained commercially and incubated at 37.8 ℃ with 60%-70% relative humidity. Embryos were staged according to the Hamburger-Hamilton system.

### In vitro virus infection and drug treatment

Cells were infected with PEDV (MOI = 1) or SARS-CoV-2 (MOI = 0.1) for 2 h at 37 ℃. After virus adsorption, inocula were removed and cells were maintained in infection medium supplemented with ferric ammonium citrate (FAC), Dp44mT, DPTP-FC, remdesivir or vehicle controls as specified. Cells were harvested at indicated time points for downstream analyses.

### In vivo virus infection and drug treatment

Animals were randomly assigned to experimental groups. Piglets, mice or chicken eggs were inoculated with PEDV, PDCoV by oral gavage or with SARS-CoV-2, IBV by intranasal administration at the indicated doses. For drug intervention, FAC or DPTP-FC was administered intramuscularly at specified doses and schedules. Body weight, clinical signs and survival were monitored daily. Animals were humanely euthanized at predetermined endpoints, and tissues were collected for histological and molecular analyses.

### Quantification of intracellular iron

Intracellular ferrous iron levels were quantified using an Iron Content Assay Kit (Elabscience, E-BC-K881-M) according to the manufacturer’s instructions.

### RNA extraction and quantitative PCR

Total RNA was extracted using TriQuick reagent (Solarbio, R1100). Reverse transcription was performed using Evo M-MLV RT Mix (Accurate Biology, AG11728). Quantitative PCR was carried out using SYBR Green Master Mix (Applied Biosystems, 4367659) on a real-time PCR system. Relative gene expression was calculated using the 2^−ΔΔCt^ method with GAPDH as the internal reference^53^.The relevant primer sequences are provided in Supplementary Table 1.

### Western blot analysis

Cells were lysed in RIPA buffer (Solarbio, R0010) supplemented with protease inhibitors (Solarbio, P0100). Proteins were separated by SDS-PAGE, transferred to PVDF membranes (Millipore, IPVH00010), and probed with the indicated primary and HRP-conjugated secondary antibodies. Signals were detected using enhanced chemiluminescence (ECL, Oriscience, PD202).The antibodies used in this study are listed in Supplementary Table 2, with sources and identifiers indicated.

### Immunoprecipitation and immunofluorescence

For IP, clarified cell lysates were incubated with antibody-conjugated protein A/G magnetic beads (Santa Cruz, sc-2003) at 4 ℃ overnight. Immunocomplexes were washed and analysed by immunoblotting. For immunofluorescence, cells or tissue sections were fixed, permeabilized, blocked, and incubated with primary antibodies followed by fluorophore-conjugated secondary antibodies. Images were acquired using a confocal microscope under identical settings.The antibodies used in this study are listed in Supplementary Table 2, with sources and identifiers indicated.

### shRNA construction

shRNA sequences targeting the indicated genes were synthesized commercially. Annealed oligonucleotides were cloned into the pSIH1-H1-copGFP-T2A-Puro vector between the EcoRI and BamHI restriction sites. Recombinant plasmids were verified by Sanger sequencing. All shRNA sequences are listed in Supplementary Table 3.

### Hematoxylin-Eosin staining

Tissues were fixed in 4% paraformaldehyde, embedded in paraffin, sectioned, and stained with haematoxylin and eosin following standard procedures. Stained sections were imaged using a digital pathology scanning system (Leica, RM2235) under identical acquisition settings for all samples.

### Viral titration

Viral titres were determined by TCID_50_ or plaque assays on Vero or VeroE6 cells using standard methods. Titres were calculated using the Reed-Muench method^52^.

### Transmission electron microscopy

Purified viral particles were negatively stained with phosphotungstic acid and visualized using a transmission electron microscope operated at 80-120 kV.

### Anaerobic purification of protein

A C-terminal His-tagged recombinant protein was expressed in HEK293F cells using a transient transfection system (Sino Biological). Briefly, HEK293F cells (3 × 10^6^ cells/mL; total volume, 800 mL) were supplemented with L-cysteine and FeCl₃ as previously described^54^ and transfected with 400 µg pcDNA3.1-nsp12-His. Cells were harvested 96 h post-transfection, washed with ice-cold PBS and lysed under strictly anaerobic conditions in a glove box. Lysates were clarified by ultracentrifugation, and the supernatants were subjected to affinity purification using Ni–NTA resin with stepwise washing. Bound proteins were eluted with buffer containing 250 mM imidazole, concentrated to approximately 150 µM using 30 kDa molecular weight cutoff centrifugal filters, and flash-frozen under anaerobic conditions for subsequent analyses.

### UV-visible absorption spectroscopy

UV-visible absorption spectra were recorded using a Shimadzu UV-2600i spectrophotometer over a wavelength range of 260-560 nm at room temperature. Measurements were performed in quartz cuvettes with a 1 cm path length, using the corresponding buffer as a blank for baseline subtraction. Spectra were collected with a scan speed of 600 nm/min and a spectral bandwidth of 1.0 nm. Representative spectra were analysed and visualized using OriginPro software.

### RNA extension assays

As previously described^55^, recombinant PEDV RdRp complex purified under anaerobic conditions was incubated with template-primer RNA duplexes and NTPs in reaction buffer. Reactions were terminated with formamide-containing quench buffer and analysed by denaturing Urea-PAGE followed by fluorescence imaging.The template-primer RNA sequences are listed in Supplementary Table 4.

### Cell-based virus RdRp activity assay

HEK293T cells were transfected with the PEDV RdRp activity assay system and the TK-Renilla luciferase internal control plasmid in a 10:1 ratio. Cells were lysed and analyzed with a dual-luciferase reporter assay kit (Promega, E1910) according to the manufacturer’s instructions.

### Electrophoretic mobility shift assay

RNA-protein binding was performed in 10 µL reactions containing 10 nM CY5- labeled cholesterol- modified probe, 1 µg IRP1 and 1× binding buffer following the manufacturer’s protocol (Beyotime, GS606). For specificity controls, reactions were run without protein, with 100× unlabeled competitor, with a mutated probe, or with a non- cholesterol probe. After 30 min on ice, samples were loaded on a pre- chilled 6% native PAGE gel in 0.5× TBE and run at 90 V for 1 h. Gels were imaged directly in the CY5 channel (Amersham ImageQuant800).The sequences of probes used for EMSA assays are provided in Supplementary Table 5.

### Virtual Screening and Molecular Docking Analysis

Virtual screening was performed using the MTiOpenScreen web server and the Diverse Chemical Compound Collection (Diverse-lib), a curated library containing 99,288 drug-like small molecules in 3D conformations^56,57^. All compounds were docked into a predefined grid centered on Val76 of the PEDV E protein (PDB: 2MM4), which encompassed adjacent residues constituting key functional motifs. The resulting candidates were ranked based on predicted binding free energies, supplemented by analysis of specific intermolecular interactions, including hydrogen bonds and hydrophobic contacts.The screening datasets supporting this study are available in the Supplementary Data Set and Extended data Table 1.

## Data Availability

All data supporting the findings of this study are available in the paper and its Supplementary Date Set. The MTiOpenScreen diverse small-molecule library used in this study can be accessed via: https://bioserv.rpbs.univ-paris-diderot.fr/services/MTiOpenScreen/. All other data are available from the corresponding author(s) on reasonable request. Source data are provided with this paper.

### Acknowledgments

The authors thank the following: Microscopy Core Facility (MCF) and the Flow Cytometry Core Facility (FCCF) at the Engineering Research Center of Efficient New Vaccines for Animals (Ministry of Education) and the Key Laboratory of Ruminant Disease Prevention and Control (West) for providing help, services, and devices. PDB DOIs: 10.2210/pdb2mm4/pdb (CoV E) and 10.2210/pdb8urb/pdb (PEDV RdRp). AlphaFold Protein Structure Database (TP73, alphafold.ebi.ac.uk/entry/O15350). Approve numbers: IACUC2024-0732 (piglets’ experiment), and 2023-015 (mice experiment). All animal experiments with SARS-CoV-2 were performed in a biosafety level 3 facility at Shenzhen Third People’s Hospital with the assistance of Dr. Lujie Fan, according to institutional and governmental guidelines of animal welfare. Breeding and housing as well as the euthanasia of the animals are fully compliant with all China applicable laws and regulations concerning the care and use of laboratory animals, including the Regulations for the Administration of Affairs Concerning Experimental Animals and the Laboratory Animal-Guidelines for Euthanasia [GB/T 39760-2021]. Details can be found in the Materials and Methods in the Supplementary Materials.

## Author Contributions

Conceptualization: M.Z. and D.T.; Methodology and Data curation: M.Z., L.H., X.F., and B.Y.; Investigation: M.Z., L.H., X.F., G.G., Z.R., and R.T.; Resources: G.L., M.D., Y.M., N.X., H.L., H.T., X.Y., J.Z., M.S., S.Y., S.L., Q.L., J.L., B.F., Y.C., Q.Z., T.Z., L.C., X.Z., X.R., L.H., Q.D., B.L., Y.H. and D.T.; Supervision: M.Z., S.D., B.L., Y.H. and D.T.; Funding acquisition: M.Z., B.L., Y.H. and D.T.; Writing-original draft: M.Z.; Writing-review & editing: M.Z., S.D. and D.T. All authors carefully read and approved the final manuscript.

## Supplementary Data Set

The top 1,500 small-molecules compounds identified in our screening have been uploaded as Dataset 1 and are accessible at https://doi.org/10.6084/m9.figshare.31436053.

## Supplementary Information

This file contains Supplementary Table 1-5.

