## Supplementary Tables for "Coronavirus envelope protein drives iron sensing disorder by hijacking the TAp73-FDXR axis"

**Supplementary Table 1**

Primer names and corresponding nucleotide sequences (5’-3’) are provided. All primers were synthesized by a commercial supplier and validated by Sanger sequencing prior to use.

**Supplementary Table 1**

| **Name** | **Sequence** |
| --- | --- |
| QPCR SARS-CoV-2 *NP* forward | CGGAATGTCTCGCATCGGTA |
| QPCR SARS-CoV-2 *NP* reverse | GGGCAACGCTTGTGTTTCAT |
| QPCR *GAPDH* forward | ACATCATCCCTGCTTCTACTGG |
| QPCR *GAPDH* reverse | CTCGGACGCCTGCTTCAC |
| QPCR PEDV *N* forward | CTTTGGTGGTAATGTGGCTG |
| QPCR PEDV *N* reverse | TCAACAGCTGTGTCCCATTC |
| QPCR *FDXR* forward | ATCACGACCACCATGACCGACAG |
| QPCR *FDXR* reverse | GGGATACCTCCTCAGCATCCAGCT |
| ChIP *FDXR* promoter forward | TGTTCCCTCACCTATGTGAA |
| ChIP *FDXR* promoter reverse | CCCATTGGGTAATAACTACG |
| QPCR *hFDXR* forward | TGGAGAGAACGGACATCACG |
| QPCR *hFDXR* reverse | GGGCTCCCGGTAACTGAATC |
| QPCR *gFDXR* forward | TTTGTGCAGCACTCCCTCTT |
| QPCR *gFDXR* reverse | GTGATCTGGTGCCACTCCAA |
| QPCR *c-FOS* forward | GTGCCAACTTCATCCCAACG |
| QPCR *c-FOS* reverse | TGGCATGGTCTTCACGACTC |
| QPCR *TfR1* forward | TTATGCCCCTCGTGAAGCTG |
| QPCR *TfR1* reverse | AGGTCACCGAGTTTTCAGCA |
| QPCR *FPN* forward | GCTGTGGTTTCATTTCGGGA |
| QPCR *FPN* reverse | AGTTCCCTCCAAGGGTTTTGG |
| QPCR *IRP1* forward | CCATTGCCAACATGTGTCCAG |
| QPCR *IRP1* reverse | AACATTCCCACAGCCTGAAGA |
| QPCR *TAp73* forward | AGCCGGACAGCACCTATTTT |
| QPCR *TAp73* reverse | TGAACTGGGACATGACAGGC |
| QPCR IBV *N* forward | TTCAACCAGATGGGCTCCAC |
| QPCR IBV *N* reverse | TTGTCTTGGCGCAGGAGAAT |
| QPCR PDCoV *N* forward | AGCTCCCAAGCGGACTTTAC |
| QPCR PDCoV *N* reverse | GAAAGTTGCGCTCAAGGTGG |

**Supplementary Table 2**

Antibodies are listed with the corresponding reagent or resource name, source, and identifier (catalog number and/or RRID, where applicable).

**Supplementary Table 2**

| **Reagent or resource** | **Source** | **Identifier** |
| --- | --- | --- |
| Beta Actin Recombinant antibody | Proteintech | Cat# 81115-1-RR; RRID: AB_2923704 |
| Influenza A Nucleoprotein / NP Antibody | Sino Biological | Cat# 11675-T62; RRID: AB_3676512 |
| FDXR Polyclonal antibody | Proteintech | Cat# 15584-1-AP; RRID: AB_2102602 |
| p73 Recombinant Rabbit Monoclonal Antibody | HUABIO | Cat# ET1609-80; RRID: AB_3069890 |
| DDDDK-Tag Rabbit mAb | ABclonal | Cat# AE092; RRID: AB_2940847 |
| mCherry Monoclonal antibody | Proteintech | Cat# 68088-1-Ig; RRID: AB_2918825 |
| HRP-conjugated Goat Anti-Rabbit IgG(H+L) | Proteintech | Cat# SA00001-2; RRID: AB_2722564 |
| Lamin B1 Polyclonal antibody | Proteintech | Cat# 12987-1-AP; RRID: AB_2136290 |
| HSCB Polyclonal antibody | Proteintech | Cat# 15132-1-AP; RRID: AB_2878110 |
| CIAO1 Polyclonal antibody | Proteintech | Cat# 10295-1-AP; RRID: AB_2260509 |
| GRP75 Polyclonal antibody | Proteintech | Cat# 14887-1-AP; RRID: AB_2120458 |
| CD71 Polyclonal antibody | Proteintech | Cat# 10084-2-AP; RRID: AB_2240403 |
| SLC40A1/FPN1 Polyclonal antibody | Proteintech | Cat# 26601-1-AP; RRID: AB_2880571 |
| Aconitase 1 Polyclonal antibody | Proteintech | Cat# 12406-1-AP; RRID: AB_10642942 |
| HA tag Polyclonal antibody | Proteintech | Cat# 51064-2-AP; RRID: AB_11042321 |

**Supplementary Table 3**

shRNA names and corresponding oligonucleotide sequences are provided. All constructs were verified by Sanger sequencing prior to use.

**Supplementary Table 3**

| **Name** | **Sequence** |
| --- | --- |
| sh-hFDXR-1 forward | GATCCGCAGAGTCGAGTGAAGACAGTCTCGAGACTGTCTTCACTCGACTCTGCTTTTTG |
| sh-hFDXR-1 reverse | AATTCAAAAAGCAGAGTCGAGTGAAGACAGTCTCGAGACTGTCTTCACTCGACTCTGCG |
| sh-hFDXR-2 forward | GATCCGCAAGTGGCCTTCACCATTAACTCGAGTTAATGGTGAAGGCCACTTGCTTTTTG |
| sh-hFDXR-2 reverse | AATTCAAAAAGCAAGTGGCCTTCACCATTAACTCGAGTTAATGGTGAAGGCCACTTGCG |
| sh-hFDXR-3 forward | GATCCGCTCAGCAGCATTGGGTATAACTCGAGTTATACCCAATGCTGCTGAGCTTTTTG |
| sh-hFDXR-3 reverse | AATTCAAAAAGCTCAGCAGCATTGGGTATAACTCGAGTTATACCCAATGCTGCTGAGCG |

**Supplementary Table 4**

List of Template-primer RNA sequences for polymerase assays used in this work.

**Supplementary Table 4**

| **Name** | **Sequence** |
| --- | --- |
| Primer strand | FAM-UUUGCAUAAUAGAUCUGUCU |
| Template strand | UUUUUUUUUUAGACAGAUCUAUUAUGCAAA |

**Supplementary Table 5**

List of probe sequences for EMSA assays used in this work.

**Supplementary Table 5**

| **Name** | **Sequence** |
| --- | --- |
| Nucleic acid aptamer | AACUUCAGCUACAGUGUUAGCUAAGUUU |
| Mutated nucleic acid aptamer | AACUUCAGCUACAAGUUUAGCUAAGUUU |
